# Hachiman is a genome integrity sensor

**DOI:** 10.1101/2024.02.29.582594

**Authors:** Owen T. Tuck, Benjamin A. Adler, Emily G. Armbruster, Arushi Lahiri, Jason J. Hu, Julia Zhou, Joe Pogliano, Jennifer A. Doudna

## Abstract

Hachiman is a broad-spectrum antiphage defense system of unknown function. We show here that Hachiman comprises a heterodimeric nuclease-helicase complex, HamAB. HamA, previously a protein of unknown function, is the effector nuclease. HamB is the sensor helicase. HamB constrains HamA activity during surveillance of intact dsDNA. When the HamAB complex detects DNA damage, HamB helicase activity liberates HamA, unleashing nuclease activity. Hachiman activation degrades all DNA in the cell, creating ‘phantom’ cells devoid of both phage and host DNA. We demonstrate Hachiman activation in the absence of phage by treatment with DNA-damaging agents, suggesting that Hachiman responds to aberrant DNA states. Phylogenetic similarities between the Hachiman helicase and eukaryotic enzymes suggest this bacterial immune system has been repurposed for diverse functions across all domains of life.

## Introduction

Helicases participate in innate and adaptive immune systems across all domains of life by sensing pathogen-associated molecular patterns (PAMPs)^1–13^. Many newly discovered antiviral defense systems in prokaryotes encode helicases homologous to diverse immune and regulatory helicases in eukaryotes (**Fig. 1A,B**)^14–16^. One such system is Hachiman, a two-gene locus encoding HamA (a protein of unknown function, DUF1837) and the Ski2-like superfamily 2 (SF2) helicase HamB. Although present in >5% of prokaryotic genomes and capable of robust protection against phylogenetically distinct phages^14,17^, molecular mechanisms governing Hachiman and many related helicase-containing immune systems remain unknown.

**Figure 1.**
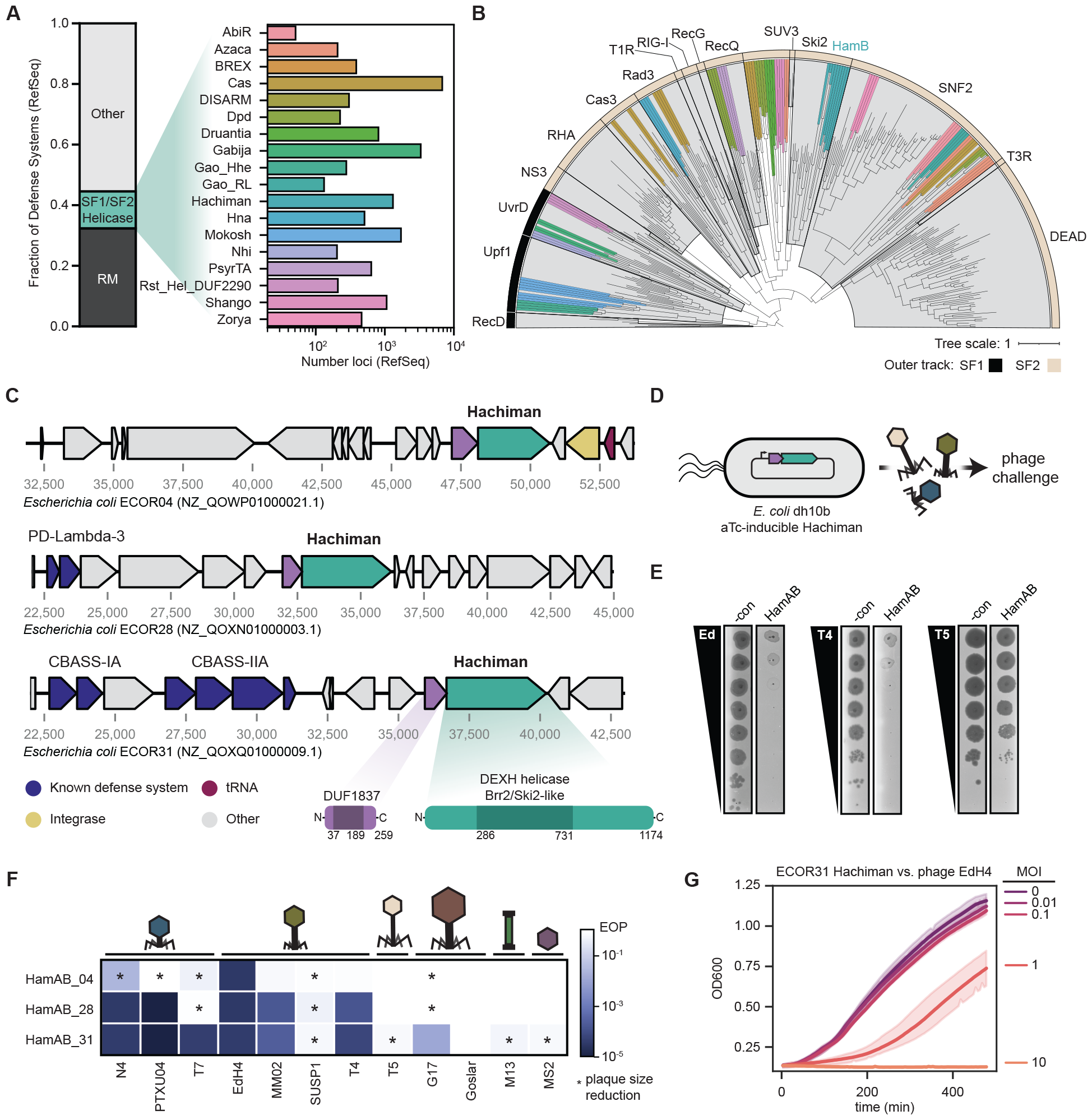
Hachiman is a two-component defense system that protects against diverse bacteriophages. **(A)** Overview of SF1/SF2 helicase-containing phage defense systems found in RefSeq genomes in the DefenseFinder database^17^. **(B)** Phylogenetic tree of core helicase domains of 329 helicases from defense systems from (A) and representative SF1/SF2 helicases^12^. Helicase superfamily is provided in the outer track (SF1 in black, SF2 in tan) and representative families demarcated in gray clades with labels. Defense system-associated helicases are colored as shown in (A). Details on tree construction and sequence alignment provided in methods. **(C)** Hachiman loci from *E. coli* strains ECOR04, ECOR28 and ECOR31 tested in this study. HamA genes are shown in purple and HamB genes shown in green. Additional defense systems identified in PADLOC^22^ are shown in blue, integrases in yellow and tRNA genes in red. All other genes are shown in gray. **(D)** Overview of phage-defense assays. Native Hachiman loci are cloned under an anhydrotetracycline (aTc)-inducible promoter, pTet, and monitored for protection against diverse phages. **(E)** Representative plaque assays for ECOR31 HamAB against sensitive phages EdH4 and T4 as well as resistant phage T5. **(F)** Comparison of different Hachiman loci against 12 diverse phages representing 12 unique phage genera. Plaque assays without EOP reductions, but a measurable difference in plaque size are denoted with an asterisk. **(G)** Protection against phage EdH4 is complete at low MOI, but insufficient at high MOI. For panels (E-G), Hachiman is induced at 20nM aTc and dCas13d targeting RFP is provided as a negative control. All assays consist of 3 biological replicates.

Here we show that despite its homology to RNA helicases, HamB is a DNA helicase that activates DNase activity of HamA upon detection of damaged DNA. Cryo-EM structures show how the HamAB complex binds DNA in different modes to enable HamA activation. Helicase ‘ratcheting’ by HamB upon substrate recognition disrupts the HamAB interface, leading to HamA release and indiscriminate degradation of host and phage DNA. Fluorescence microscopy shows that Hachiman clears host and phage DNA simultaneously. DNA-depleted cells remain intact, while cells lacking Hachiman succumb to phage-induced lysis. The observation of Hachiman activation in the absence of bacteriophage but in the presence of DNA-damaging agents suggests that Hachiman responds to DNA damage that accumulates during cell stress. Biochemical and structural data imply ATP-bound HamAB contacts intact DNA, enabling detection of genome integrity and activation by HamA release when DNA damage surpasses a normal threshold. HamA nuclease activity likely creates additional sites for Hachiman binding and activation, leading to amplification of the immune signal and culminating in broad restriction of phylogenetically diverse phages.

HamB helicase domain organization and its ability to regulate the HamA effector enables controlled activation that may be a principle of other helicase-containing defense pathways. Like Hachiman, other defense systems may act in response to cell stressors including but not limited to phage infection.

## Results

### Hachiman confers broad-spectrum protection against diverse bacteriophages

Helicases are common components of immune systems in eukaryotes^9,18^. This is also true for prokaryotic immune systems. Of the ∽150 defense systems cataloged in DefenseFinder^17^, 18 contain an SF1/SF2 helicase, comprising nearly 20% of non-restriction-modification (RM) defense loci identified in RefSeq (**Fig. 1A**)^19^. We sought to put these common proteins in the context of known helicases. Using 95 representative defense system helicases and 236 well-characterized representative SF1/SF2 helicases^12^, we performed a phylogenetic analysis of the core helicase domain (**Fig. 1B**, see Methods). We confidently assigned 20 of the 25 helicases to an established helicase subfamily, spanning 7 subfamilies: SF1_UvrD (GajB, Nhi, Rst_DUF2290 Helicase), SF1_Upf1 (Hhe, MkoA, MkoC), SF2_Cas3 (Cas3), SF2_Rad3 (Csf4, Hna), SF2_RecQ (DpdF, PsyrT), SF2_Ski2 (HamB) and SF2_SNF2 (AbiRc, BrxHII, DpdE, DrmD, Gao_RL, ZacC, ZorD). The Hachiman-encoded HamB protein is closely related to SF2 Ski2 helicases, orthologs of which have diverse activities on RNA and ssDNA substrates in eukaryotes and archaea (**Fig. 1B, Fig. S1**)^20,21^.

To establish a cell-based assay for assessing Hachiman function, we first identified three different Hachiman loci in *E. coli* strains ECOR04, ECOR28 and ECOR31 using PADLOC (**Fig. 1C**)^22^. We expressed these three Hachiman loci under aTc-inducible control in *E. coli* and challenged the cells in plaque assays using *E. coli* phages representing twelve distinct genera (**Fig. 1D-E, Tables S2-S4**)^23^. HamAB from ECOR31 conferred the greatest degree of defense, providing 10^2^-10^5^-fold reduction in efficiency of plaquing for eight diverse dsDNA phages (**Fig. 1F**), consistent with the broad-spectrum activity of *Bacillus cereus* Hachiman against *Bacillus subtilis* phages^14^. Hachiman conferred near-complete defense against sensitive phages at low multiplicity of infection (MOI<1), but diminished defense at high viral doses (MOI>1) (**Fig. 1G**), consistent with an abortive infection (Abi) phenotype in which infected cells die before phage infection matures, preventing viral spread^24^.

The four phages resistant to the three tested Hachiman systems (T5, MS2, M13 and Goslar) possess unique genome properties within this phage panel. MS2 is an ssRNA phage and lacks a dsDNA genome, while M13 uses rolling-circle replication to produce its ssDNA genome^25^. The dsDNA genomes of T5 and Goslar have limited accessibility to defense systems as they are compartmentalized before and during infection, respectively^26–28^. Overall, phage challenge experiments suggest Hachiman activity protects against diverse dsDNA phages by recognizing and subverting a central feature of dsDNA phage infection.

### Structural basis of HamAB complexation

To determine the molecular basis for Hachiman function, we purified HamA and HamB individually. Only HamB was soluble in isolation. Coexpression of the complete native Hachiman locus produced a complex of HamA and HamB (**Fig. S3A-B**). A 2.7 Å cryo-EM structure of the HamAB complex is a 1:1 heterodimer (HamA1:HamB1, **Fig. 2A, Fig. S3A-G, Table S1**). The domain organization of HamB is generally consistent with Ski2/Brr2 helicases, with two stacked RecA-like helicase domains, RecA1 and RecA2, comprising the helicase core^29^. A degenerated winged-helix domain (WH*) and C-terminal α-helical region (CAH, C α-helix) form the likely nucleic acid binding cleft (**Fig. 2B**). At the N-terminus, an α-helical bundle (NAH, N α-helical) common to HamB orthologs but not found in related Ski2 helicases, contributes to binding HamA. At the C-terminus, a barrel-like fold reminiscent of oligonucleotide binding (OB) domains sits on the side of the complex. HamB folds with intact helicase motifs, including active site DEGH resides in the helicase core (**Fig. S3H-J**).

**Figure 2.**
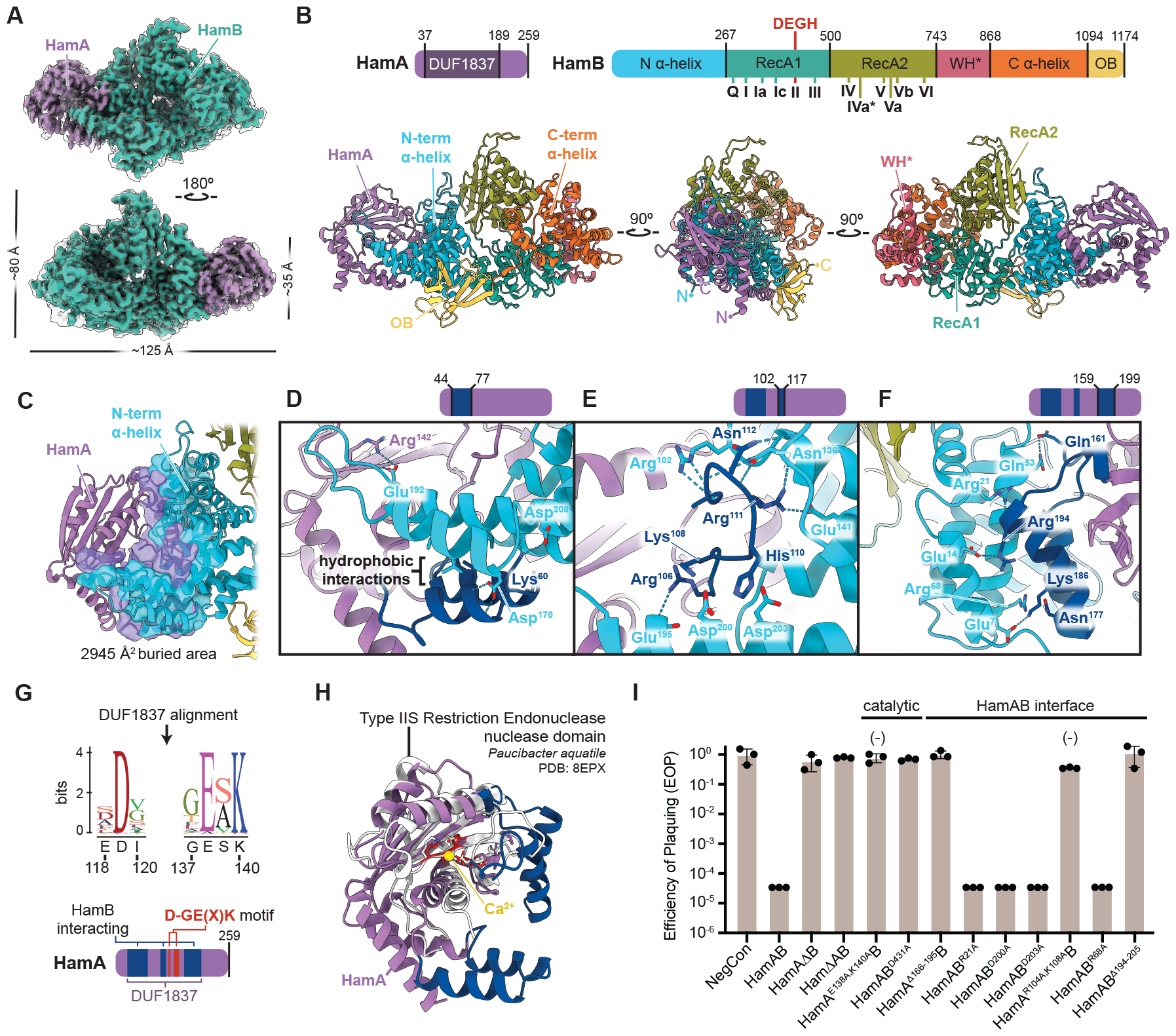
Structural basis of Hachiman complexation and identification of the HamA active site. **(A)** Cryo-EM density of the *E. coli* ECOR31 apo HamAB complex. The sharpened map is colored, while the unsharpened map is overlaid and transparent. **(B)** Orthogonal views of the HamAB structure, with domains colored according to the key above. Walker motifs are annotated in the HamB RecA1 and RecA2 domains. **(C)** Overview of the HamA-HamB NAH interface, with surfaces involved in the interaction shown. **(D-F)** Detail of three subregions, HamA^44-77^ (D), HamA^102-117^ (E), and HamA^159-199^ (F), contributing to the AB interface. Residues contributing to hydrogen bonding interactions are shown as sticks and are labeled with colors corresponding to the key above each view and in (B). **(G)** Sequence logo resulting from alignment of HamA DUF1837 ORFs. The ECOR31 HamA sequence and corresponding position is shown below each residue logo. **(I)** Plaque assays demonstrating the ability of HamAB and various mutants to confer defense against phage EdH4. Individual data points of three independent biological replications are shown along with the mean and standard deviation. The (-) symbol indicates a reduction in plaque size.

The apo HamAB structure shows how HamA contacts the HamB NAH, with three AB interface regions contributing to 2,945 Å^2^ of total buried surface area (**Fig. 2C**). The first subregion contains a helix-loop-helix which stacks with a HamA helix-loop-helix in the reverse orientation (HamA^44-77^, **Fig. 2D**). The second region is in the center of the interface and includes numerous hydrogen bonding contacts between HamB and an extended HamA-interacting loop (HamA^102-117^, **Fig. 2E**). The third subregion on the HamB N-terminal side is another instance of helical docking with predicted hydrogen bonding and nonpolar interactions (HamA^159-199^, **Fig. 2F**). Structural analysis of the AB interface helps explain why HamA was insoluble in isolation and suggests that HamAB may assemble cotranslationally in the native cellular context (**Fig. 2D**).

### HamA DUF1837 encodes a nuclease

HamA binds HamB, but its role in immunity is unknown. Alignment of HamA sequences characteristic of DUF1837 revealed a highly conserved D-GE(X)K motif, consistent with a metal ion-dependent enzyme capable of phosphodiester hydrolysis (**Fig. 2G**)^30^. Structurally, HamA is most similar to the Type IIS restriction endonuclease from *Paucibacter aquatile*^*31*^, with conservation of the core helix/sheet motif (**Fig. 2H, Fig. 2SK**). HamA diverges from the *P. aquatile* nuclease in regions of DUF1837 that facilitate binding to HamB (**Fig. 2C-G**).

Cell-based phage defense assays showed that deletion of HamA or HamB, or mutation of their putative active site residues to create HamA^E138A,K140A^B (HamA*B, nuclease-deficient) or HamAB^D431A^ (HamAB*, helicase-deficient), ablated defense (**Fig. 2I**). Although single interface mutations failed to impact defense, double mutation of a conserved interface motif (R/K)XX(R/K) or deletion of entire helix-loop-helix motifs in HamA blocked Hachiman function. Together, these results show that Hachiman requires both nuclease (HamA) and helicase (HamB) activities for function, and that complexation of HamA and HamB is essential for phage defense.

### HamB is a DNA helicase

Despite its functional requirement for phage defense, the identity of the HamB helicase substrate is unclear. Using a malachite green assay that detects orthophosphate release during NTP hydrolysis, we found strong HamB ATPase activity in the presence of single-stranded DNA (ssDNA, **Fig. 3A**). To test HamB DNA helicase activity, we performed DNA unwinding assays by incubating HamB with DNA duplexes of varying lengths and single-stranded overhangs. HamB was capable of ATP-dependent unwinding of a 15 bp duplex with a 15 nt 3′ overhang (**Fig. 3B**). In addition, HamB unwinds forked, 5′ overhang and blunt DNA duplexes, albeit with lower efficiency compared to 3′ overhang-containing substrates, suggesting promiscuous substrate acceptance (**Fig. 3C-E, Fig. S4A-F**). Longer duplex lengths are not well tolerated (**Fig. 3F**). Testing of different DNA duplex and overhang lengths showed that HamB can process a range of DNA substrates, and that it prefers longer 3′ overhangs (**Fig. S4A-F**).

**Figure 3.**
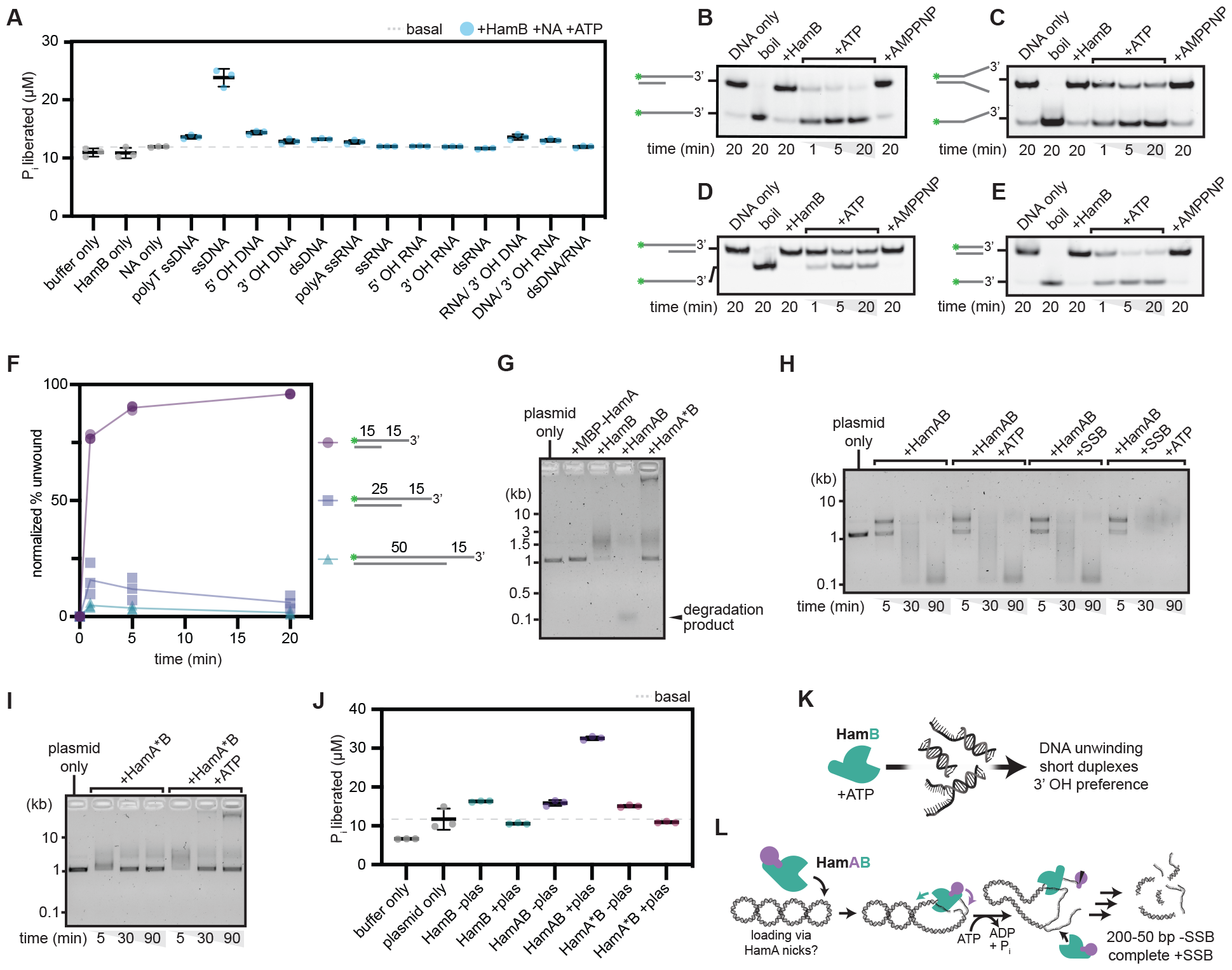
HamAB is a DNA nuclease/helicase that degrades plasmids in vitro. **(A)** Malachite green ATPase assays of HamB against a panel of nucleic acid substrates. Individual data points of three independent biological replicates and the mean and standard deviation are shown. **(B-E)** HamB DNA unwinding assays on substrates with a 15 bp duplex and a 15 nt 3′ OH (B), forked 15 nt OH (C), 15 nt 5′ OH (D), and no overhang (E). DNA substrates are labeled with 5′ FAM. Gels are representative of three independent replicates. **(F)** Normalized percent unwinding of DNA substrates with 15 bp (circles), 25 bp (squares), and 50 bp (triangles) duplex lengths, all labeled with 5′ FAM and with a 15 nt 3′ OH. Individual data points shown are quantifications of replications of unwinding assays in the format of B-E normalized against basal unwinding (see Methods). **(G)** *In vitro* plasmid clearance assay after 90 min at 37ºC with ATP using MBP-HamA, HamB, HamAB, and HamA*B visualized on a 0.75% agarose gel. **(H)** Time course of HamAB plasmid clearance with addition of ATP or SSB visualized on a native agarose gel. **(I)** Time course assay as in J with mutant HamA*B. **(J)** ATPase activity of HamB, HamAB, and HamA*B with or without supercoiled plasmid substrates. Individual data points of three independent biological replicates and the mean and standard deviation are shown. **(K)** Cartoon summarizing *in vitro* activities of HamB. **(L)** Cartoon depicting a model for HamAB-mediated plasmid degradation.

### HamAB degrades plasmids *in vitro*

HamB unwinds DNA substrates, and HamA is a putative nuclease. To determine whether HamA cuts DNA and to ascertain the combined functions of the HamAB complex, we tested Hachiman activity against purified plasmid DNA. Titration of the wildtype HamAB complex, but not HamA*B, in reactions with supercoiled plasmid DNA show initial plasmid nicking followed by a ladder of degradation products (**Fig. 3G, Fig. S4G**). Degradation products converged to sizes between 50 and 200 bp. Supporting our identification of the DUF1837 active site, this observation is consistent with HamA acting as a nuclease effector in Hachiman immunity (**Fig. 2G-I**). Although HamA*B cannot cleave DNA, it forms a low-mobility species upon addition of ATP that may represent a different binding mode captured only when the HamA nuclease is catalytically deactivated.

To assess the possible influence of phage-encoded single-stranded binding (SSB) protein, which has been implicated in activating Hachiman and other phage defense systems^32–34^, we incubated reactions with either *E. coli* SSB (*Ec*SSB) or phage T4 SSB prior to addition of Hachiman components. Both types of SSB induced HamAB- and ATP-dependent plasmid degradation, arguing against direct recognition of phage SSB in Hachiman activation (**Fig. 3H, Fig. S4I-L**). In time-course experiments, Hachiman degrades plasmid DNA within ten minutes when *Ec*SSB is present, amounting to an order of magnitude rate increase (**Fig. S4K-L**). *Ec*SSB potentially mediates ssDNA accessibility by prevention of reannealing (**Fig. 3H**)^35^. *Ec*SSB did not have an observable effect in HamA*B time courses (**Fig. S4J**), suggesting that HamA nuclease activity is required for subsequent DNA unwinding *in vitro* (**Fig. 3I**).

We noted that the low mobility species observed in HamA*B-plasmid reactions accumulates in an ATP-dependent manner (**Fig. 3I**). Furthermore, ATPase assays with HamB, HamAB and HamA*B in the presence of plasmid DNA reveal that unlike HamAB, both HamB and HamA*B suppress ATPase activity upon substrate addition (**Fig. 3J**). The low mobility species may therefore represent a state in which HamA*B ATP-binding enables association with, but not cleavage of, intact dsDNA. By extension, these data imply not only that plasmid destruction requires DNA cleavage by HamA, but also that this activity may be coupled with HamB ATPase activity (**Fig. 3J**). Considering that HamAB does not require ATP to degrade plasmid DNA in the absence of SSB, we postulate that HamAB loads DNA ends induced by HamA nicks *in vitro*, triggering ATP hydrolysis which in turn activates HamA-mediated degradation (**Fig. 3L**). When the HamA nuclease is inactivated, HamB does not load DNA ends, but can nonetheless bind intact DNA in an ATP-dependent manner.

### Structural basis of HamB DNA binding

Our biochemical studies suggest two possible modes of DNA binding, one that triggers ATP hydrolysis, and one that enables HamAB binding to intact dsDNA (**Fig. 3J**). We observed ATPase activity upon incubation of HamB with a mixed base ssDNA substrate. We performed cryo-EM analysis on a complex of HamB and ssDNA with ATP added during complexation. In the resulting 2.8 Å reconstruction, HamB retains the general domain organization observed in the apo HamAB structure (**Fig. 4A-B, Fig. S5A-I, Table S1**), but lacks the extended helix-loop-helix domain structure that contributes to HamA binding (**Fig. 2D, Fig. 4B**). Disorder of this region in the HamB-DNA structure further supports its important role in complexation. Surprisingly, despite the addition of only ssDNA to HamB during sample preparation, we observed duplex DNA in the cryo-EM density. The duplex, which is 8 bp in length, arises from a partially palindromic region of the DNA substrate. The duplex appears partially unwound, with one 3’ end bound within the HamB nucleic acid binding pocket (**Fig. 4C**). Several residues including, but not limited to, canonical Walker motifs contribute to ssDNA binding in the helicase core (**Fig. 4D, Fig. S3H-I**). In the midsection of the duplex, a ‘pin’ reminiscent of the strand unwinding wedge in the PriA primosomal helicase lies in the center of the duplex and pries the strands apart by pi stacking and physical occlusion (**Fig. S5J**)^36,37^. Observation of a 3’ end in the entry site of the helicase is consistent with the 3’ OH preference determined from *in vitro* assays (**Fig. 3B**). There is no obvious site for the 3’ end of the ssDNA molecule to exit the DNA pocket (**Fig. 4C**).

**Figure 4.**
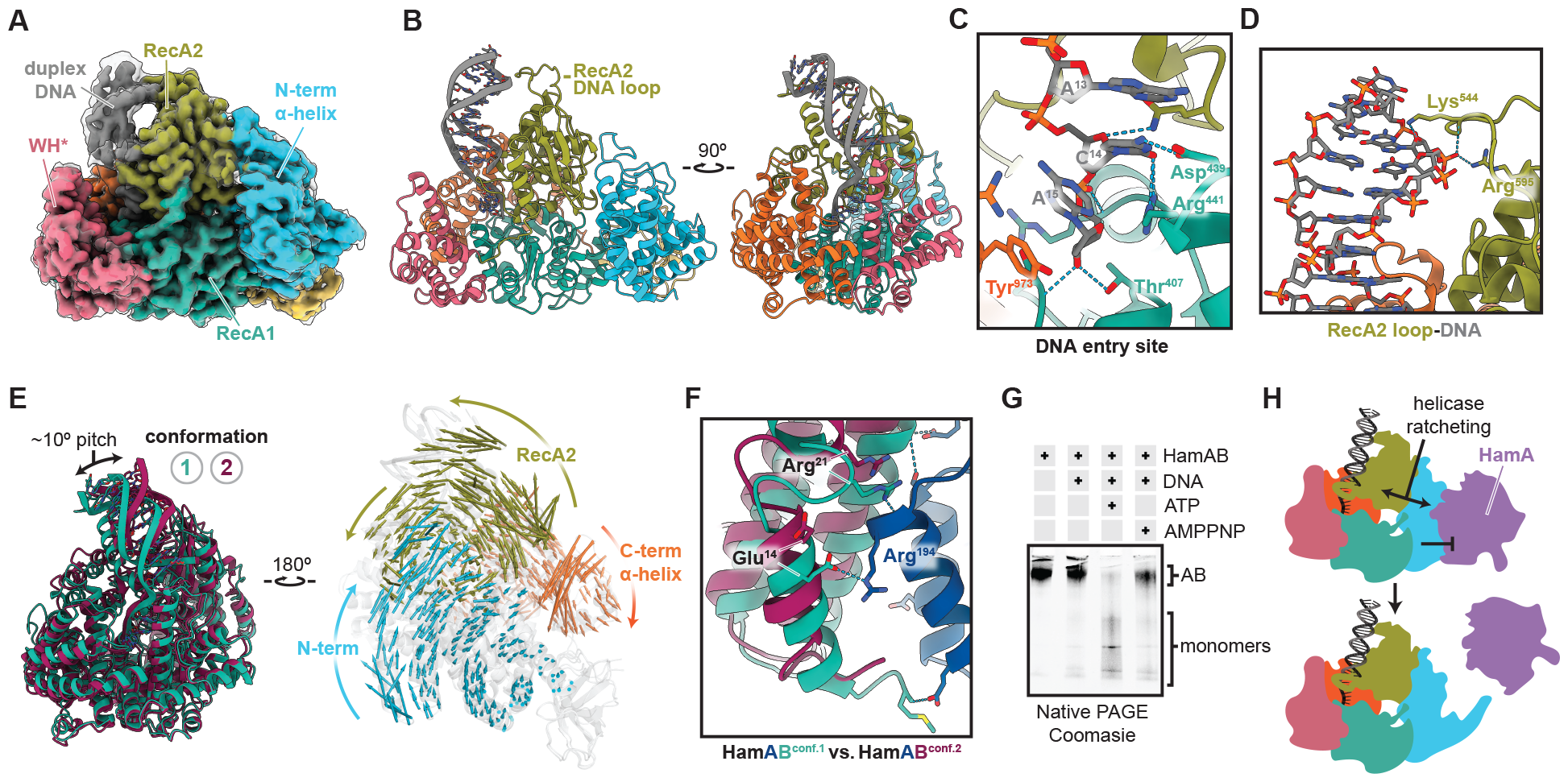
Structural basis of HamB-DNA binding and helicase ratcheting. **(A)** Cryo-EM density of the 2.8 Å HamB-DNA density. The sharpened map is colored according to domain, while the unsharpened map is overlaid and transparent. **(B)** Orthogonal views of the 2.8 Å HamB-DNA structure. **(C)** Detail of the 3’ end of the DNA buried within the DNA entry site of HamB. Hydrogen bonds and contributing residues are shown with a dashed line. **(D)** Detail of the DNA duplex-interacting RecA2 loop. **(E)** Left, superimposed conformers of HamB-DNA viewed from the DNA side, with conformation 1 (2.8 Å) colored teal and conformation 2 (2.9 Å) colored burgundy. Right, conformations 1 and 2 viewed from the NAH side and transparent, with vectors colored according to domain representing motion between the two conformations. Vectors are scaled 2x and are calculated using modevectors. **(F)** Representative disruption of the predicted AB interface between the two HamB conformations. AB interactions disrupted by HamB motion are shown and labeled. **(G)** Native PAGE of reactions of the HamAB complex with the DNA where ratcheting was observed in cryo-EM. ATP and DNA appear to dissociate the AB complex. **(H)** Model for HamB signal transduction to the NAH and concomitant release of HamA.

### Helicase ratcheting may release HamA

We noticed significant conformational variability in the HamB-DNA particle ensemble. Using 3D variability analysis and 3D classifications, we resolved an alternative conformation (conformation 2) of HamB to a nominal resolution of 2.9 Å (**Fig. S5A-I, Table S1**, Methods). In the alternate conformation, we observe significant repositioning of the RecA2, NAH and CAH domains coupled with pitching of the DNA duplex by approximately 10º (**Fig. 4E**, left). DNA contacts and the RecA1, WH* and OB folds remain virtually unchanged. Viewed from the HamA direction, the NAH and CAH rotate clockwise, while RecA2 moves counterclockwise (**Fig. 4E**, right). Movement of the NAH:RecA2 interface is especially dramatic and involves remodeling of interface regions. For example, Tyr^525^ and Trp^521^ residues on a distal RecA2 sheet contacting the NAH shift ∽9 Å between the two conformations (**Fig. S5K-L**). We propose that dynamic switching between HamB conformations represents helicase ‘ratcheting’ upon entry of the DNA substrate into the active site and triggering of ATPase activity. Motion of the RecA2 domain upon helicase ratcheting transduces to the NAH (**Fig. 4E**). The NAH is responsible for binding the HamA nuclease (**Fig. 2C-F, 2I**).

We superimposed each conformation of HamB bound to DNA with HamB in the apo complex structure. In conformation 1, HamB is in approximately the same position as HamB in the AB complex. Conversely, changes in conformation 2 appear to disrupt predicted interactions with HamA (**Fig. 4F**), including hydrogen bonds in HamA^159-199^, a region shown to be essential in phage challenge assays (**Fig. 2I**).

Structural evidence suggests helicase ‘ratcheting’ transduces motion to the NAH which disrupts the AB interface. To test this prediction, we incubated HamAB with the mixed base ssDNA seen in HamB-DNA structures. When ATP and DNA are present, the AB complex dissociates into monomers (**Fig. 4G**). AMPPNP appears to block dissociation, suggesting ATP hydrolysis provides input energy to trigger conformational switching which disrupts the AB interface. Together, these data support a model in which structural changes in HamB release the HamA nuclease, which then degrades DNA (**Fig. 4H**). These changes appear to activate upon entrance of ssDNA into the active site of the helicase. In further support of this model, biochemical evidence from plasmid assays demonstrate that nicked DNA triggers ATPase activity, while intact DNA untouched by HamA nucleolytic activity does not activate ATP hydrolysis (**Fig. 3J**).

### Hachiman degrades phage and host DNA simultaneously

We propose that Hachiman binds and degrades DNA. To connect the proposed HamAB structural states with cellular activities, we visualized Hachiman responding to phage infection *in vivo* using fluorescence microscopy. In uninfected cells, expression of either WT or nuclease-deficient HamAB does not impact normal cellular activities. Hachiman does not affect nucleoid morphology, consistent with our observation of little or no toxicity upon expression of Hachiman in growth experiments. When we challenged cells with sensitive phage EdH4 (**Fig. 1F**), we observed that, throughout the course of the infection, HamAB-bearing cells were completely devoid of DAPI signal (**Fig. 5A,B**). These ‘phantom’ cells occur at a higher frequency in strains harboring normally functioning Hachiman. When HamA*B is present, or if the helicase Walker B motif is deactivated in the HamAB* mutant, phantom cells form less frequently (**Fig. 5A,B**). Instead, the cytoplasm appears to be filled with decondensed DNA . Some cells contain numerous DNA-rich puncta indicative of phage packaging prior to lysis (**Fig. 5C**). The vast majority of uninfected cells have intact chromosomes for all three strains. 60 minutes post infection with EdH4, 36.5% of cells expressing WT HamAB have mostly or fully degraded host chromosomes (**Fig. 5B**). However, the fraction of cells that lack visible DNA was reduced for strains expressing inactivated HamAB mutants. These observations are consistent with an abortive infection phenotype and agree with biochemical and structural data identifying Hachiman as a DNA-degrading defense system.

**Figure 5.**
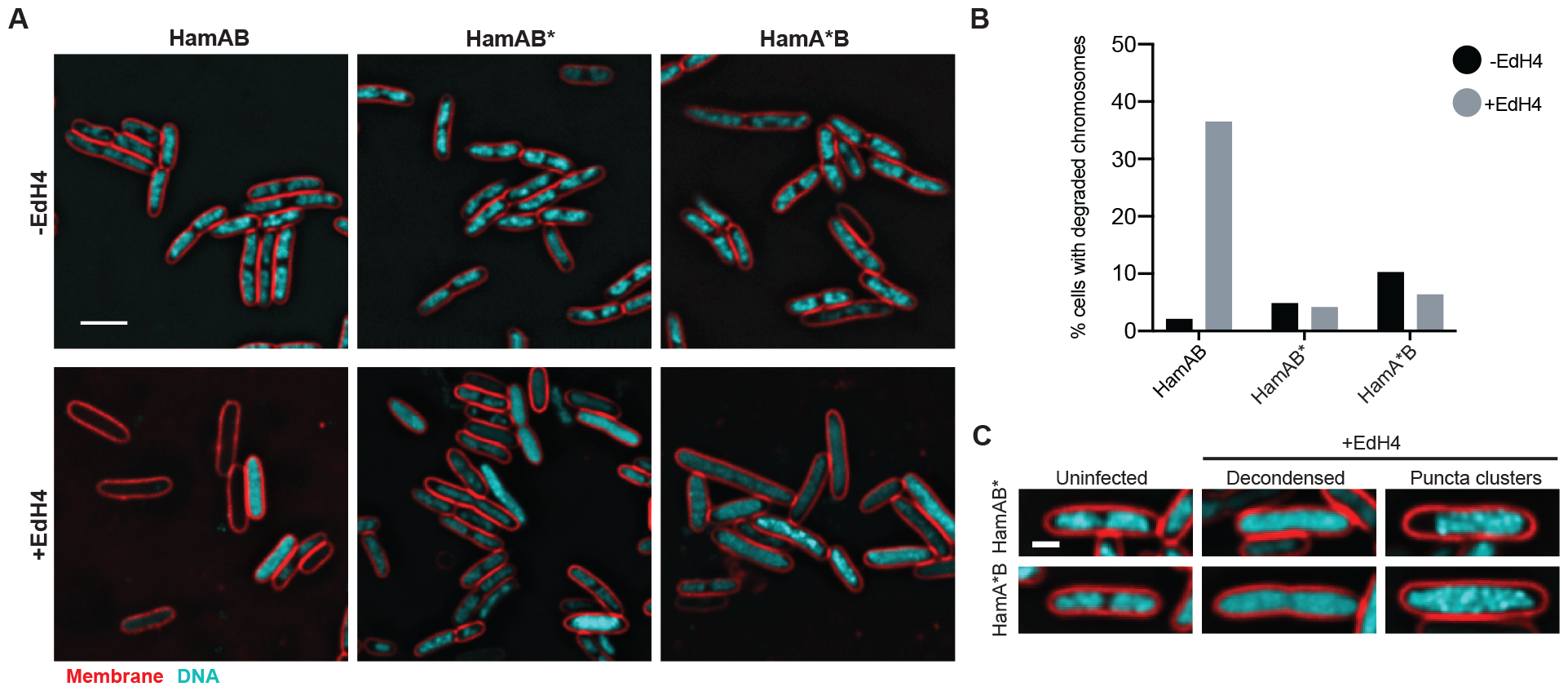
Hachiman defends against bacteriophage and prevents lysis by nonspecific DNA clearance. **(A)** Wildtype HamAB, HamAB* (HamAB^D431A^) or HamA*B (HamA^E138A,K140A^B) expressing *E. coli* either uninfected (-EdH4) or 60 minutes post-infection by phage EdH4 (+EdH4). Scale bar = 3 µm. Cell membranes were stained with FM4-64 (red)^38^ and DNA was stained with DAPI (cyan). **(B)** Quantification of percentage of cells with completely degraded chromosomes. The following lists the sample size, in number of cells, for each condition: WT HamAB, -EdH4 = 188 cells; WT HamAB, +EdH4 = 211 cells; HamAB*, -EdH4 = 185 cells; HamAB*, +EdH4 = 213 cells; HamA*B, -EdH4 = 171 cells; HamA*B, +EdH4 = 180 cells. **(C)** Examples of DNA morphologies in uninfected and EdH4-infected cells expressing HamAB* or HamA*B. Scale bar = 1 µm. Cell membranes were stained with FM4-64 (red)^38^ and DNA was stained with DAPI (cyan).

### DNA damage activates Hachiman

Hachiman responds to phage infection by clearing cells of DNA. The HamB helicase recognizes DNA ends (**Fig. 2B-F**), leading to ATP hydrolysis (**Fig. 2J**), which in turn releases the HamA nuclease (**Fig. 4E-H**). Since Hachiman is capable of defense against diverse bacteriophage genera with little or no protein homology, we considered the possibility that Hachiman does not recognize a conserved phage component such as phage-encoded SSB^32^. Instead, Hachiman could directly sense general changes in host physiology via general DNA damage.

We reasoned that small molecules that interfere with DNA metabolism might differentially engage Hachiman if it indeed responds to changes in host genome integrity. To test this hypothesis, we treated cells with subinhibitory amounts of nalidixic acid (Nal), a quinolone inhibitor of DNA gyrase and topoisomerase IV (topo IV), essential enzymes that regulate DNA supercoiling and catenation^39,40^. Nal intercalates DNA in the normally transient protein-linked cleavage complex generated during strand passage in gyrase and topo IV, locking them in the open form^41–44^. Aberrant persistence of the protein-DNA linkage results in DNA nicks, replication fork arrest, and double-strand breaks (DSBs). In the absence of bacteriophage, we observed growth inhibition in response to subinhibitory amounts of nalidixic acid when WT Hachiman was present compared to inactivated mutants (**Fig. 6A, Fig. S6A**). Next, we treated cells with novobiocin (novo), an aminocoumarin that also interferes with gyrase and topo IV, but by an orthogonal mechanism - novo competes with ATP for binding in a distal pocket, resulting in loss of ATPase activity^45,46^. Because novo does not directly interfere with strand passage, it does not cause direct DNA damage. In growth experiments, WT and mutant HamAB had no differential effects on growth after novo treatment (**Fig. 6B, Fig. S6B**). Likewise, cultures treated with gentamycin (gm)^47^, an aminoglycoside that inhibits translation by binding to the 30S ribosomal subunit, responded equally to the drug for all Hachiman constructs (**Fig. 6C, Fig. S6C**). Our results demonstrate that Hachiman can be triggered in the absence of bacteriophage, that activation is the result of DNA damage and that this activation requires the combined catalytic activities of HamA and HamB.

**Figure 6.**
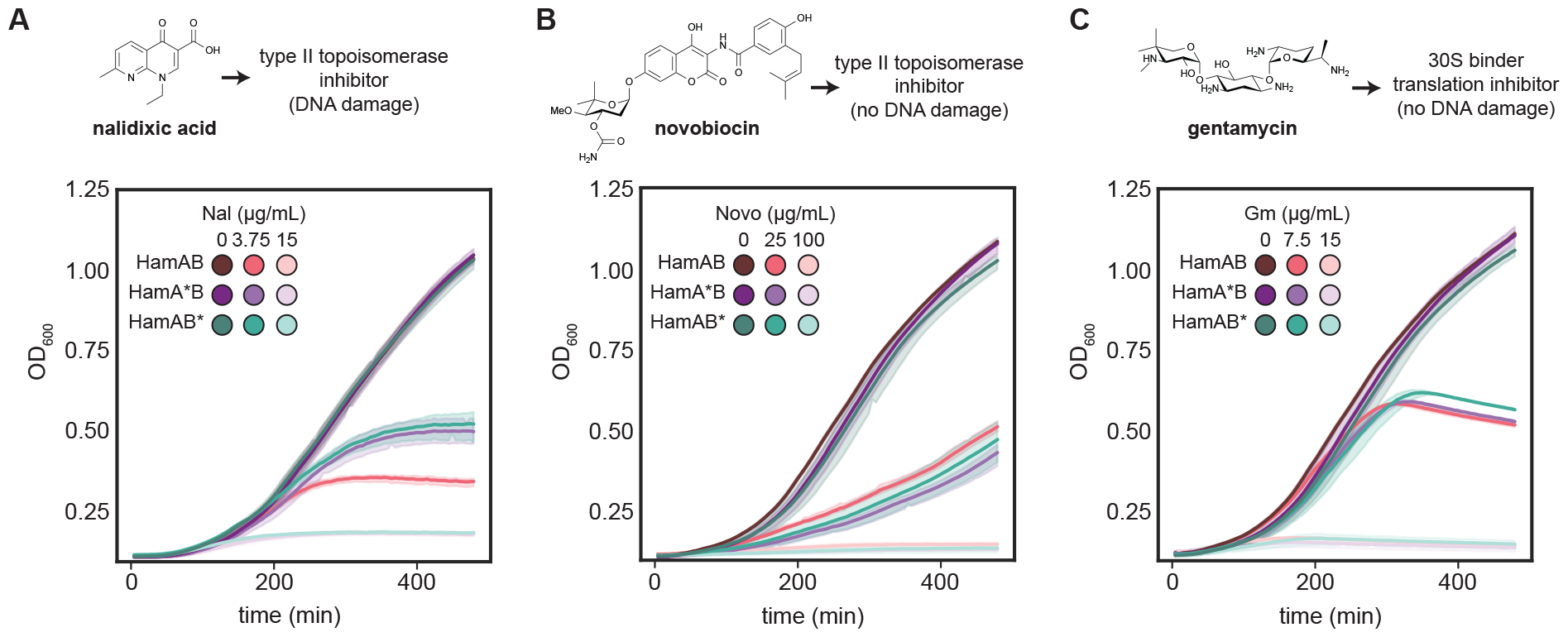
DNA damage activates Hachiman. **(A-C)** Cell growth of *E. coli* expressing wildtype HamAB, HamA*B and HamAB* at 20nM aTc at 20nM aTc in the absence or presence of subinhibitory and inhibitory concentrations of nalidixic acid (A), novobiocin (B) or gentamycin (C). Growth curves are colored according to condition. See Fig. S6 for complete minimum inhibitory concentration (MIC) determinations. All measurements were performed in biological triplicate.

### Hachiman scans intact dsDNA

Our data are consistent with a model in which Hachiman triggers abortive infection when ssDNA enters the HamB active site, triggering ATP hydrolysis and HamA release. Although the exact mechanism of DNA damage ‘sensing’ by HamB remains unclear, we observed formation of ATP-dependent HamA*B-DNA complexes *in vitro* (**Fig. 3I**). We used cryo-EM to visualize this state. HamA*B was incubated with plasmid DNA and ATP for 30 minutes of reaction, after which the specimen was frozen (**Fig. 7A**). In the resulting micrographs, many particles can be seen binding to intact plasmid DNA (**Fig. 7B**). In 2D class averages of plasmid-bound particles, complete HamA*B complexes are seen, with duplex DNA spanning the protein and bending slightly at the point of contact (**Fig. 7C, Fig. S7A**). The angle of the DNA in this ‘scanning’ state is orthogonal to DNA resolved in the ‘loading state’ in HamB-DNA structures (**Fig. 7D**). Masked 3D classification and unbiased alignments produced a map with a 3.2 Å nominal resolution, with lower resolutions (5–7 Å) for the plasmid DNA, although the major and minor grooves in the central region are apparent (**Fig. 7E,F, Fig. S7A-E, Table S1**). The dsDNA interacts with the RecA2 loop region (**Fig. 7E**). There are few differences between the rest of the complex and the apo HamAB structure (**Fig. 2B**). In the molecular model, duplex DNA occupies the same location as RecA2 loop - we could not find an alternate conformation of the loop structure, leading us to conclude that it becomes disordered once DNA is bound, opening to accommodate the duplex (**Fig. 7F**). ATP occupies the binding pocket, consistent with biochemical results (**Fig. 3I-J**). Contacts made with ATP are in agreement with predictions for HamB Walker and helicase motifs (**Fig. 7G, Fig. S3H-I**). Our observations suggest that HamAB surveys DNA in an alternative ‘scanning’ mode which enables traversal across intact dsDNA. In this mode, which is facilitated by the RecA2 DNA loop, DNA is restrained from the nuclease active site. HamAB could conceivably activate if a ssDNA end enters the active site either during surveillance of dsDNA or, alternatively, by loading of a DNA end (**Fig. 7H**). Entrance of ssDNA into the helicase active site triggers ATPase activity, leading to structural rearrangements enabling release of HamA.

**Figure 7.**
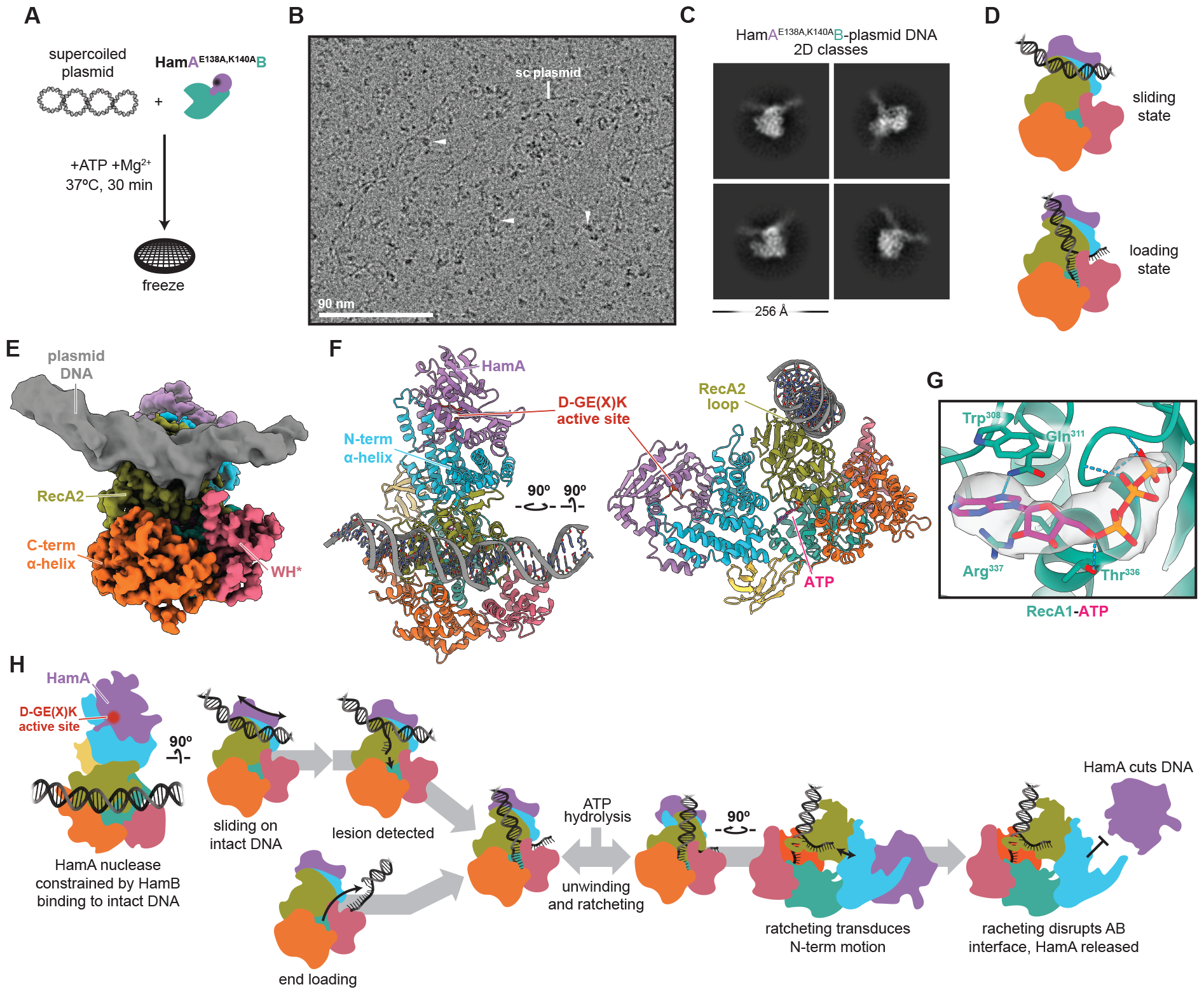
Hachiman scans intact dsDNA. **(A)** HamA*B-plasmid +ATP specimen preparation. **(B)** Representative motion-corrected, dose-weighted cryo-EM micrograph from the HamA*B-plasmid DNA dataset. Plasmid DNA and bound particles are indicated with white arrows. **(C)** Representative 2D classes of particles bound to plasmid DNA. **(D)** Cartoon depicting the scanning state resolved here and comparison with the loading state resolved in the HamB-DNA dataset. **(E)** Composite cryo-EM density colored according to domain. Protein regions are from the 3.2 Å deepEMhancer-sharpened map, while DNA is from the 3.2 Å sharp map masked and B-factor refined to display helical features. **(F)** Orthogonal views of the HamA*B-plasmid DNA structure. The DNA sequence is unknown. **(G)** Detail of ATP in HamB, with residues and hydrogen bonds shown. The density is masked to ATP. **(H)** Proposed mechanism of Hachiman immunity.

## Discussion

Our results reveal that the Hachiman prokaryotic defense system is a nuclease-helicase complex, HamAB, that responds to changes in genome integrity. Upon contact with a free ssDNA end, the end inserts into the HamB active site to induce ATP hydrolysis and consequent release of the HamA nuclease. The released HamA catalyzes rapid DNA degradation, creating intact ‘phantom’ cells cleared of both phage and host DNA. The total DNA degradation phenotype is reminiscent of NucC-mediated clearing in Type III CRISPR-Cas systems^48^. That Hachiman separates the nuclease and helicase components may be intrinsically linked to its robust Abi phenotype, as *trans* nuclease activity could initiate a positive feedback loop. Elevated DNA damage activity from HamA would enable other HamAB complexes to load new sites of DNA damage, amplifying the immune response and DNA clearance. This model explains how Hachiman avoids self-targeting: during normal cell activities, DNA damage is limited and transient due to proofreading activities of the native DNA repair machinery. We propose that major changes in genome integrity, such as host-genome degradation or, potentially, recombination-dependent replication encoded by dsDNA phages^25^, result in accumulated damage that ‘tips the scales’, leading to HamAB activation during infection. Thus, cells harboring Hachiman wield broad-spectrum defense, but at the risk of autoimmunity.

The spread of ‘selfish’ mobile genetic elements such as phage entails substantial changes to nucleic acid metabolism in the cell. For example, phage T4 is known to degrade the host genome to preferentially replicate and suppress antiphage activities^49^, while also encoding programs to interfere with host transcription^50^. In addition to host interference, lytic phages engage in rapid replication and lysis. Viral teleonomy entails tradeoffs. Faster replication results in higher error rates^51^, leading to more lesions and stalled replication forks in addition to damage to the phage genome induced by other defense systems^52^. Phages are known to engage in orthogonal homologous recombination to compensate, a process that universally involves a free 3’ ssDNA end and displacement-loop intermediates^53,54^. We propose that Hachiman senses and activates in response to universal motifs associated with stress on the integrity of host or phage DNA^55^, enabling infection sensing across a range of infection strategies. This explains the broad spectrum protection conferred by Hachiman defense. Whether Hachiman acts on plasmid invasion and replication intermediates is unclear. We imagine that Hachiman is evolutionarily tuned to activate only when DNA damage is too severe for the host DNA repair machinery to remedy or if invading genomes undergo uncontrolled replication^25,53^.

We identify HamA, previously DUF1837, as the effector nuclease responsible for DNA clearance. Compared with structural homologs in Type IIS restriction-modification systems, HamA contains insertions that mediate interactions with HamB. Considering the HamA interaction domain (NAH) is present in all HamBs, and that this interaction domain is absent in close relatives in the Ski2 subfamily (**Fig. S5M**)^56,57^, we propose HamA insertions were acquired during Hachiman evolution to enable nuclease regulation and allosteric activation. Another distinguishing feature of HamB is the loop on the crown of the RecA2 domain which both facilitates ‘scanning’ of intact dsDNA and forms contacts with DNA during loading into the helicase active site. Comparison to predicted structures of HamB orthologs confirms that either the RecA2 loop or a highly positively charged patch exists at this position, suggesting dsDNA sliding or binding may be a common feature in Hachiman defense (**Fig. S7G**). Other immune helicases have been proposed to ‘scan’ DNA or RNA for pathogenic signatures^13,58^. Future studies could address with single molecule techniques whether HamB-DNA association via the RecA2 loop confers diffusive or unidirectional motion to aid in genome integrity sensing.

Ski2-like helicases variably accept RNA or DNA substrates^29^. Our biochemical results demonstrate HamB is more functionally similar to the Ski2-like DNA helicase Hel308 than Ski2 RNA helicases involved in RNA regulatory processes such as splicing and mRNA decay. Hel308 is conserved in archaea and metazoans, but is absent in bacteria and fungi^59,60^. Like HamB, Hel308 has a wide substrate scope, with a preference for 3’ to 5’ DNA unwinding^20,57^. Intriguingly, Hel308 was shown to localize to sites of DNA damage induced by the topoisomerase I inhibitor camptothecin in human cells^61^. We observed an analogous response to DNA damage induced by the Type II topoisomerase inhibitor nalidixic acid in cells harboring functional Hachiman. In addition to genetic similarity, functional symmetries between HamB and Hel308 suggest a close evolutionary relationship.

The AbpAB antiphage defense system was described before the identification of Hachiman^62^. AbpAB encodes a nuclease (AbpA) and a Ski2-like helicase (AbpB) with combined activity against DNA. The N-terminal domain of AbpA is similar to the Cap4 endonuclease domain from CBASS systems^63^. The C-terminal domain of AbpA is remarkably genetically and structurally similar to HamA, but contains catalytically inactive residues within the HamA catalytic site (**Fig. 2, Fig. S2A, Fig. S7H**). AbpAB was recently shown in cell-based assays to activate in response to mitomycin C-induced DNA damage, in agreement with our data^34^. Based on structural and functional homology, we propose AbpAB is a Hachiman variant with an N-terminal fusion in the HamA homolog AbpA (**Fig. S2A,B**). Why AbpAB would encode a catalytically inactive form of the HamA nuclease while carrying an additional, distinct endonuclease remains enigmatic.

A recent study proposed phage-encoded SSB activates *B. cereus* Hachiman^32^. However, our data suggest this mode of activation is a proxy for the true activator of Hachiman encoded in DNA structure. One parsimonious explanation is that expression of phage SSB is incompatible with host replication and recombination machinery while the phage preferentially replicates its own genome. Potentially, this could result in DNA. *In vitro* activation of ECOR31 HamAB activity did not require SSB, nor was a differential effect seen when comparing phage and host SSB, consistent with observations in the XPD SF2 helicase^64^. Considering the minimal sequence and structural homology between the two stimulatory SSBs, we consider it unlikely that either activates HamB by protein-protein interactions. Our results imply other defense systems believed to be SSB-activated may be stimulated by DNA damage^34,65^.

This study provides the first structural and biochemical analyses of Hachiman function, extending our understanding of mechanisms of prokaryotic immunity. The Hachiman sensor helicase HamB has surprising functional similarities to the archaeal and metazoan DNA repair helicase Hel308, which may explain its general activity against DNA damage. Genome integrity sensing may be a more general role of helicases in immune systems beyond Hachiman.

## Acknowledgements

The authors thank members of the Doudna Laboratory for discussions and reading of this manuscript. We thank D. Toso and R. Thakkar of the Cal-Cryo facility @QB3 Berkeley for assistance in cryo-EM data acquisition. This work was supported by the United States National Institutes of Health (NIH U19AI135990, NIH U01AI142817), the United States National Science Foundation (NSF 1817593) and the Howard Hughes Medical Institute (HHMI). B.A.A. was supported by m-CAFEs Microbial Community Analysis & Functional Evaluation in Soils (m-CAFEs{at}lbl.gov), a Science Focus Area led by Lawrence Berkeley National Laboratory based upon work supported by the US Department of Energy, Office of Science, Office of Biological & Environmental Research under contract number DE-AC02-05CH11231. E.G.A. and J.P. were funded by the Howard Hughes Medical Institute Emergent Pathogens Initiative grant and National Institutes of Health grants R01-GM129245. E.G.A. was previously supported by NIH PiBS training grant (T32 grant GM133351). This paper was typeset with a bioRxiv template by @Chrelli: www.github.com/chrelli/bioRxiv-word-tem-plate.

## Author contributions

O.T.T. and B.A.A. conceived of this project. B.A.A. performed phylogenetic analyses with assistance from O.T.T.. B.A.A. and A.L. performed molecular cloning, phage challenge assays and pharmacological assays. O.T.T. and J.J.H. performed protein purification and biochemical experiments with assistance from J.Z.. Cryo-EM data were collected by O.T.T. Cryo-EM data processing was performed by O.T.T. and J.J.H.. E.G.A. performed live cell microscopy experiments. This work was supervised by J.A.D. and J.P.. The manuscript was written by O.T.T., B.A.A., and J.A.D.. All authors contributed to reviewing and editing this manuscript.

## Competing interest statement

J.A.D. is a co-founder of Caribou Biosciences, Editas Medicine, Intellia Therapeutics, Mammoth Biosciences and Scribe Therapeutics, and a director of Altos, Johnson & Johnson and Tempus. J.A.D. is a scientific advisor to Caribou Biosciences, Intellia Therapeutics, Mammoth Biosciences, Inari, Scribe Therapeutics, Felix Biosciences and Algen. J.A.D. also serves as Chief Science Advisor to Sixth Street and a Scientific Advisory Board member at The Column Group. J.A.D. conducts academic research projects sponsored by Roche and Apple Tree Partners. J.P. has an equity interest in Linnaeus Bioscience Incorporated and receives income. The terms of this arrangement have been reviewed and approved by the University of California, San Diego, in accordance with its conflict-of-interest policies.

## Data availability

Structure coordinates and corresponding density maps have been deposited at the Protein Data Bank (PDB) and Electron Microscopy Database (EMD), respectively, under the following accessions: *E. coli* ECOR31 apo HamAB PDB 8VX9, EMD-43613; *E. coli* ECOR31 HamB-DNA (conformation 1) PDB 8VXA, EMD-43615; *E. coli* ECOR31 HamB-DNA (conformation 2) PDB 8VXC, EMD-43616; *E. coli* ECOR31 HamA(E138A,K140A)B-plasmid DNA PDB 8VXY, EMD-43643. This paper does not report original code. Plasmids used will be deposited on Addgene upon publication of this manuscript. Additional materials and information are available from the lead contact upon request.

## Materials and Methods

### Helicase and Hachiman phylogenetic analysis

Proteins chosen for phylogenetic analysis were from DefenseFinder RefSeq db with a RefSeq Protein ID^16,17^. Proteins from non-Restriction-Modification (RM) defense systems encoding a SF1/SF2 helicase domain were further selected for phylogenetic comparison: AbiR (AbiRc), Azaca (ZacC), BREX (BrxHI, BrxHII), DISARM (DrmA, DrmD), Dpd (DpdE, DpdF, DpdJ), Druantia (DruE), Gabija (GajB), Gao_RL (RL), Hachiman (HamB), Hhe (HheA), Hna (Hna), Mokosh (MkoA, MkoC), Nhi (Nhi), PsyrTA (PsyrT), Rst Helicase+DUF2290 (Helicase), Shango (SngC), Type I CRISPR-Cas (Cas3), Type IV CRISPR-Cas (Csf4/DinG), and Zorya (ZorD). Helicase proteins without RefSeq protein IDs or not tracked in DefenseFinder (ex. Hma) and from RM defense systems (ex. Type I (Type_II_REases) and Type III RM (Type_III_REases)) were not included in the analysis.

For analysis of SF1 and SF2 helicases, 4 randomly chosen examples of the above defense-associated SF1/SF2 helicases were selected and compared to a curated set of SF1/SF2 helicase core domains ^12^. To focus our analysis on the helicase core domain, we manually curated the helicase core domain of defense-associated helicases through structural alignment followed by sequence alignment. First, a predicted AlphaFold2 structure of each defense-associated helicase was aligned to the core helicase domain of HamB (PDB ID: 8VXA, this work, residues 286-472, 500-731)^17,66,67^ within the core helicase domain were inferred. Next, for each helicase type, representative helicases were aligned to the corresponding annotated reference using MUSCLE (default parameters), core helicase annotation extracted and insertions removed in Geneious Prime v2023.2.1 ^68,69^To build the helicase tree, sequences were concatenated with a curated set of SF1/SF2 core helicase domains^12^, aligned using ClustalOmega (default parameters), phylogenized using IQ-TREE (-bb 1000, -m MFP (optimal model: LG+R8)), bootstraps inferred using UFBoot2 and visualized using iTOL^70–73^. Clades were inferred by using bootstrap values ≥ 85 followed by analysis of associated sequences.

For analysis of full-length HamA and HamB, proteins were combined and clustered using mmseqs2 (80% coverage, 80% sequence identity)^74^. HamA nucleases and HamB helicases investigated in this study from ECOR04, ECOR28, and ECOR31 and from prior work^14,34^ were spiked into this collection of non-redundant protein sequences for in-group analysis. Sequences were aligned with ClustalOmega (default parameters), phylogenized using IQ-TREE (-bb 1000, -m MFP (optimal model: LG+F+R8 (HamA), LG+F+R10 (HamB))) and visualized using iTOL^70,71,73^. Clades were inferred by using bootstrap values ≥ 85 in the HamB followed by analysis of spiked-in in-groups.

### Identification and visualization of Hachiman-containing loci

Hachiman loci and nearby defense systems were identified in ECOR04 (NZ_QOWP01000021), ECOR28 (NZ_QOXN01000003) and ECOR31 (NZ_QOXQ01000009) using PADLOC^22^. Gene annotations were visualized and represented using dna_features_viewer^75^.

### Plasmid and strain construction

All new plasmids in this study were constructed through PCR, gel extraction (Zymo D4001) and Gibson assembly^76^. DNA PCR templates for wildtype and mutant Hachiman loci originated from isolated gDNA (Qiagen DNeasy Blood & Tissue Kit, 69504) from ECOR04, ECOR28 and ECOR31. For all *E.coli* assays, Hachiman loci were cloned under pTet control into a p15a vector with Cm resistance. For protein expression and purification, Hachiman loci were cloned under T7 control in a high copy vector with Carbenicillin resistance. In general, plasmids were propagated in dh10b genotype *E.coli* (F – mcrA Δ(mrr-hsdRMS-mcrBC) endA1 recA1 φ80dlacZΔM15 ΔlacX74 araD139 Δ(ara, leu)7697 galU galK rpsL (StrR) nupG λ-) (Intact Genomics). For subsequent phage assays (see plaque assays), some plasmids were transferred to *E.coli* MC1000, *E.coli* ECOR47 or *E.coli* dh5a F’ (NEB, C2992). For protein expression and purification, plasmids were transformed into BL21 AI genotype *E.coli* (F-ompT hsdSB (rB-mB-) gal dcm araB::T7RNAPtetA).

Plasmids used in this study are listed in Table S3. All plasmids used in this study were sequenced-confirmed by full-plasmid sequencing services using Primordium.

### Phage propagation and scaling

All phage propagation was performed using commonly employed protocols. In general, phages were propagated at 37°C in LB Lennox media using an initial MOI of 0.1 and host *E.coli* BW25113 (F-DE(araD-araB)567 lacZ4787(del)::rrnB-3 LAM-rph-1 DE(rhaD-rhaB)568 hsdR514). Phage G17 was propagated on *E.coli* DSM 103255. Phage Goslar was propagated on *E.coli* MC1000. Phages MS2 and M13 were propagated on *E.coli* dh5a F’ cells with added 1mM CaCl_2_. All phage titers were determined on their assay hosts harboring a negative control plasmid (pBA635).

### Plaque assays

Phage plaque assays were performed using a double agar overlay protocol. Briefly, cultures were grown overnight at 37 °C and 250 rpm. To form overlays, 100 µL of saturated culture was mixed with molten LB Lennox agar (0.7% w/v agar, 60°C). For assays involving G17 or Goslar a less-dense agar concentration was used (0.35% w/v). The agar-bacterial mixture was supplemented with Cm to a final overlay concentration of 34 µg/mL and anhydrotetracycline (aTc) (Sigma) concentration of 20 nM. For phages M13 and MS2 an CaCl_2_ was added to a final concentration of 1 mM. The top agar and bacterial mixture was poured onto a 5 mL LB Agar and Cm plate and left to dry under microbiological flame for 15 minutes. Phages were diluted 10X in SM buffer (Teknova) and 2µL of each dilution were spotted onto the top agar and left to dry under microbiological flame. Once dry, plates were incubated at 30°C for 12-16 hours. Plates were scanned in a standard photo scanner and plaque forming units (p.f.u) were enumerated, keeping note of changes in plaque size relative to a negative control. During assays where “lysis from without^77^” phenotypes were observed, we interpreted these as a lack of productive phage infection and were approximated as 1. p.f.u. at that concentration. Efficiency of plaquing (EOP) calculations were calculated as mean(p.f.u.condition)/mean(p.f.u.negativecontrol) in Python. All plaque assays were performed in biological triplicate. Visualizations were performed using GraphPad Prism or Seaborn in Python.

### Growth assays

Liquid phage experiments were performed in a Biotek plate reader using LB +Cm +20 nM aTc media. Strains containing a Hachiman-expressing plasmid or negative control (pBA635) were grown overnight at 37 °C and 250 rpm. Strains were seeded into a 96-well microplate reader plate (Corning 3903) at a cfu of ∽8e6 cfu per well. For phage experiments, phages were diluted 10X in SM buffer (Teknova) and 4 µL added to each well to achieve MOI = 10 (or 8e7 pfu phages) in the maximal case. For antibiotic (Nalidixic acid (Sigma), novobiocin (Sigma), and gentamycin (Sigma)) experiments, antibiotics were diluted 2X in LB + Cm media and 4µL added to each well to achieve final, maximal concentrations of 30 µg/mL, 100 µg/mL, or 15 µg/mL respectively. Growth was monitored in a Biotek Cytation 5 plate reader for 16 hours at 800 rpm shaking at 37°C with OD600 readings every 5 minutes. All assays were performed in biological triplicate, sourcing strains from independent overnights. Data were plotted using the seaborn package in Python. Subinhibitory concentrations of antibiotic were determined by investigating concentrations of antibiotic that consistently grew to a lower carrying capacity than a no-drug control, but higher than the maximal concentration of antibiotic.

### DNA Substrate Preparation

Oligonucleotides were synthesized by Integrated DNA Technologies (Coralville, IA). Substrates used in unwinding assays were prepared by mixing the fluorescent or larger strand with a 1.5-fold excess of the non-fluorescent strand in hybridization buffer (20 mM Tris-HCl (pH 7.5), 25mM KCl, 10mM MgCl_2_), and heating to 95 °C followed by slow cooling to room temperature for at least an hour. Annealed substrates were purified on an 8% native PAGE gel.

### Protein expression and purification

All Hachiman purification constructs were N-terminally tagged with 10xHis-MBP-TEV. For complex purification vectors in the native locus format, only HamA was tagged with 10xHis-MBP-TEV. After transformation into BL21-AI *E. coli*, cells were grown to an optical density of ∽0.6 then induced overnight at 16ºC with 0.5 mM isopropyl-β-D-thiogalactopyranoside (IPTG) and 0.1% L-arabinose. Cells were harvested and resuspended in lysis buffer (20 mM HEPES, pH 8, 500 mM NaCl, 10 mM imidazole, 0.1% Triton X-100, 1 mM Tris, 1 mM (2-carboxyethyl)phosphine (TCEP), 1 tablet crushed complete EDTA (ethylenediaminetetraacetic acid)-free protease inhibitor (Roche), 0.5 mM phenylmethylsulfonyl fluoride (PMSF), and 10% glycerol). Cells were lysed by sonication, then clarified by centrifugation. The clarified lysate was incubated with Ni-NTA resin for 1 hr. The resin was washed with wash buffer (20 mM HEPES, pH 8, KCl mM NaCl, 10 mM imidazole, 1 mM TCEP, and 5% glycerol), then bound protein was eluted with wash buffer supplemented with 300 mM imidazole. Eluate was then run over an MBPTrap column (GE Healthcare), washed with MBP/SEC wash buffer (20 mM HEPES, pH 8, 150 mM KCl, 1 mM TCEP, and 5% glycerol), and eluted with MBP/SEC buffer supplemented with 10 mM maltose. Eluted protein from the MBPTrap column was treated with TEV protease overnight. Protease-treated samples were concentrated and run on either a Superdex 200 10/300 GL column (Cytiva) for HamA or HamB solo constructs, or a Superose 6 increase 10/300 (Cytiva) for HamAB complex preparations. Aliquots were snap-frozen in liquid nitrogen for later use.

### Cryo-EM sample preparation and data acquisition

The apo HamAB complex sample was rerun over a Superose 6 increase 10/300 (Cytiva) column in Cryo-EM buffer (20 mM HEPES, pH 8,100 mM KCl, 1 mM TCEP, and 0.5% glycerol). The HamB-DNA complex sample was prepared by combining 15 µM HamB with 20 µM DNA in Cryo-EM buffer supplemented with 1 mM ATP and 2 mM MgCl_2_ and reacting for 30 min at room temperature. Samples were then purified over a Superdex 200 10/300 GL column (Cytiva) in Cryo-EM buffer. The HamA*B-plasmid DNA sample was prepared by combining HamA*B with 1 µg plasmid in cryo-EM buffer supplemented with 1 mM ATP and 2 mM MgCl_2_. The reaction was incubated at 37ºC for 5 min, after which the sample was frozen. Samples were frozen using a FEI Vitrobot Mark IV cooled to 8 ºC at 100% humidity on 2/2 200 mesh UltrAuFoil gold grids (Electron Microscopy Sciences) glow discharged at 15 mA for 25 s (PELCO easyGLOW). In all cases, 4 ul of each specimen was applied to the grid and immediately blotted for 5 s with a blot force of 8 units.

For apo HamAB and HamA*B-plasmid datasets, micrographs were collected on a Titan Krios G3 equipped with a GATAN K3 Direct Electron Detector in CDS mode and a BIO Quantum energy filter operated at 300 kV and 81,000x nominal magnification in super-resolution mode (0.465 Å/pix). For the HamB-DNA dataset, micrographs were collected on a Talos Arctica equipped with GATAN K3 Direct Electron Detector operated at 200 kV and x36,000 magnification in super-resolution. All cryo-EM data was collected using SerialEM v.3.8.7 software. Images were obtained in a series of exposures generated by the microscope stage and beam shifts. For the HamAB apo and HamA*B-plasmid datasets, movies were acquired in an 11×11 pattern. For the HamB-DNA dataset, movies were acquired in a 7×7 pattern.

### Cryo-EM data processing

All Cryo-EM data processing was performed in cryoSPARC (v4.2.0 or v4.3.0)^78^. For the HamAB apo specimen, 4,796 movies were collected and 2× binned to a calibrated pixel size of 1.05 Å. 3,314 exposures were accepted after patch motion correction and patch contrast transfer functions (CTF). First, 5,538,550 particles from blob picking were subjected to 2D classification and *ab initio* reconstruction of 3 classes, yielding a density consistent with a complete heterodimeric AB complex. The initial *ab initio* volume was used to create 100 evenly-spaced projection-based templates for further template picking. The 3,505,636 particles from template picking were subjected to 4 class *ab initio* reconstruction, which gave a HamAB density of 865,990 particles. Further 2D classification and 2D rebalancing (with 9 superclasses) were used to mitigate orientation bias and remove rod-shaped particles missing multiple HamB domains, leading to a final set of 309,630 particles. Single-class *ab initio* reconstruction followed by non-uniform refinement with on-the-fly defocus and CTF refinement steps gave the final 2.65 Å map^79^, which was sharpened using DeepEMhancer^80^.

For the HamB-DNA specimen, 9,133 movies were collected and 2× binned to a calibrated pixel size of 1.12 Å. A total of 8,906 exposures were accepted after patch motion correction and patch CTF. Template picker using templates generated from the HamB AF2 prediction gave the best results and were used to isolate 18,254,957 particles at a box size of 256 pix. Reasonable 2D classes were used to train deep picker, which was used to infer 1,125,114 particles at a larger box (512 pix). *Ab initio* reconstruction followed by non-uniform refinement gave a consensus 2.76 Å density with considerable heterogeneity. Then, 3D Variability Analysis (3DVA) with 3 modes using the ‘simple’ output was used to visualize continuous motion presented in Supplementary Movie 1^81^. 3D classification with 5 classes was used to resolve densities representing the maxima of motion resolved in 3DVA. Class 1 of the 3D classification gave HamB-DNA confirmation 1, which was refined (non-uniform refinement with on-the-fly defocus and CTF optimization) to 2.79 Å and sharpened with DeepEMhancer. Class 0 was refined and sharpened in the same manner, giving HamB-DNA conformation 2 at 2.93 Å.

For the dataset containing HamA*B incubated with plasmid DNA, 3724 movies were corrected for beam-induced motion using patch motion correction, then 2× binned to a calibrated pixel size of 1.05 Å. Contrast transfer function parameters were calculated using patch CTF. Initially, 16,398,369 particles were picked using blob picker from all 3724 micrographs. Multiple rounds of reference-free 2D classification were subsequently performed to remove “bad” particles (i.e., particles in 2D classes with fuzzy or uninterpretable features) yielding 87,084 particles with clear protein characteristics. The particles were then submitted for Topaz training, and the resulting Topaz model was used to pick particles from all 3724 micrographs^82^, giving a total of 1,322,669 particles. Multiple rounds of reference-free 2D classification were subsequently performed to remove junk particles. After selecting the best classes, 317,540 particles were used for *ab initio* reconstruction of 3 classes. Of the 3 classes, 2 classes were selected for subsequent heterogeneous refinement. Heterogeneous refinement yielded a good class with 205,538 particles, and non-uniform refinement was performed with the particles from this class, yielding a reconstruction at 2.86 Å resolution. Afterward, multiple rounds of reference-free 2D classification were performed again to select for good particles which presented resolvable features from 2D classification, resulting in 103,451 particles selected. Then, *ab initio* reconstruction was performed on the selected particles, and subsequently non-uniform refinement, which resulted in a 2.96 Å reconstruction. Then, a focused 3D classification with 4 classes was performed on the predicted DNA binding region of HamA*B, based on views seen in 2D classification, to classify for DNA-bound HamA*B. To generate the focus mask, an atomic model of B-form DNA was built at the predicted DNA binding region, and then a mask of the predicted DNA binding region was artificially simulated using ChimeraX’s molmap function with subsequent binarization and softening. The solvent mask was generated to contain both the protein and predicted DNA densities. The best class containing 29,904 particles yielded a classification that was enriched for DNA-bound HamA*B. Then non-uniform refinement was performed on those particles, which resulted in a 3.2 Å reconstruction, which was then sharpened with deepEMhancer.

### Model building

The initial model of HamAB was obtained with the ColabFold^67^. To build the model, we fit the Colabfold prediction into the experimental HamAB apo density with the fitmap tool in UCSF ChimeraX v1.6.1^83^. There were significant differences in nearly every region of the structure which required iterative manual refinement with a combination of Coot v0.9.4.1^84^, ISOLDE v1.6.0^85^, and Phenix 1.20.1-4487^86^. The HamAB apo structure served as the initial model for all other models. The HamB-DNA and HamA*B-plasmid DNA models were built in the manner described above. DNA was built *de novo*. In the HamA*B-plasmid DNA dataset, the DNA sequence could not be determined, so DNA was modeled as a 31-mer of A-T to maintain base pair interactions during model building. All models were subjected to a final round of Phenix real-space refinement.

### NTPase assays

Orthophosphate liberation was determined with a Malachite Green Phosphate Assay kit (BioAssay Systems, Hayward, CA, USA) according to the manufacturer protocol. Substrate design was inspired by a recent study^87^. Briefly, HamB reactions were run in Isothermal Amplification Buffer (henceforth IAB, New England Biosciences, 20 mM Tris-HCl, 10 mM (NH_4_)_2_SO_4_, 50 mM KCl, 2 mM MgSO_4_, and 0.1% Tween® 20 (pH 8.8 at 25 °C). HamB was diluted to 40 nM, and nucleic acid substrates were diluted to 100 nM, or 4 ng/µl for plasmid reactions, in a total reaction volume of 80 ul in a clear bottom, flat, black 96-well assay plates (Corning Costar). Reactions were allowed to sit for at least 15 min at ambient temperature before initiation with addition of ATP to 1 mM and incubation at 37ºC. Reactions were quenched after 30 min with the addition of activated malachite green reagent. The absorbance values of wells were measured after 20 min of color development at ambient temperature with a Biotek plate reader at 620 nm. Orthophosphate liberation was interpolated against a standard curve with known concentrations of free phosphate.

### Plasmid activity assays

Plasmid interference assays were conducted in IAB buffer. Plasmids were diluted to 4 ng/µl, and ATP was added to a final concentration of 1 mM where indicated. In cases where *E. coli* (QIAGEN) or phage T4 SSB (gp32, New England Biosciences) were added, the final concentration was either 400 nM or 4 µM in limiting and saturating conditions, respectively, and the SSB-DNA mixture was allowed to rest for 15 min on ice. Reactions were started with addition of MBP-HamA, HamB, HamAB, or HamA*B to a final concentration of 500 nM, unless otherwise noted, and were incubated at 37ºC. Reactions were quenched with addition of EDTA to 10 mM at various time points and were imaged on 0.75% TBE agarose gels. Gels were stained with SYBR-safe and imaged on a ChemiDoc MP (BioRad).

### Gel-shift helicase unwinding assays and quantification

Unwinding reactions were carried out at 30 °C in IAB buffer. 100nM of HamB was incubated with 20 nM DNA substrate for 5 min, and the reactions with protein were initiated by addition of ATP or AMP-PNP to a final concentration of 1 mM. At either 1 min, 5 min, or 20 min, reactions were quenched on ice with STOP Buffer (0.4 U proteinase K (NEB), 18 mM EDTA, 0.36% SDS, and 9% glycerol. Boiled substrates were incubated at 95 °C for 5 minutes before immediate loading. Samples were electrophoresed until separation in an 8% TBE polyacrylamide gel at 4 °C. Fluorescent bands were imaged using a Typhoon FLA scanner and quantified using Fiji. The fraction of unwound substrate by HamB was estimated by dividing the intensity of the unwound strand over the sum of the intensities of the unreacted duplex and unwound strand, minus the fraction of unwound substrate from spontaneous unwinding without HamB at 20 minutes, then normalized to the fraction of unreacted duplex without HamB at 20 minutes:

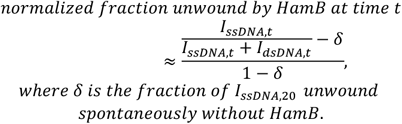

### Live single-cell static fluorescence microscopy

Fluorescence microscopy experiments were performed in at least biological duplicate. Host cells were grown to OD_600_ 0.5 in LB (+chloramphenicol 30 µg/mL) at 30 °C. 8 µL were spotted and spread on the surface of 1% agarose, 25% LB imaging pads containing 30 µg/mL chloramphenicol and 0.05 nM aTc on single-well concavity glass slides, then incubated for 1.5-2 hours at 30°C without coverslips in a humidor. At this stage, 1 µL of ∽10^11^ PFU/mL EdH4 lysate was spotted and spread onto the imaging pads and the pads were incubated at 30°C without coverslips in a humidor until the desired infection time point.

All live cell microscopy was performed on a DeltaVision Elite Deconvolution microscope (Applied Precision, Issaquah, WA, USA). Imaging pads were stained with 8 µL of dye mix (25 µg/mL DAPI, 3.75 µg/mL FM4-64) at room temperature and a glass coverslip was placed on top of the pad immediately before imaging. DAPI: Thermo Fisher Scientific Cat# D1306, FM4-64: Thermo Fisher Scientific Cat# T13320. Cells were imaged with 10-15 slices in the Z-axis at 0.2 µm increments.

Images were deconvolved in DeltaVision SoftWoRx (version 6.5.2). Image analysis was performed using raw images in FIJI (version 2.3.0/1.53q) and GraphPad Prism (version 10.0.0). Figure panels were created in Adobe Photoshop (21.2.0), GraphPad Prism (version 10.0.0), and Adobe Illustrator (24.2).

