## Supplemental Figures 1-7, Supplementary Tables 1-6 for "Hachiman is a genome integrity sensor"

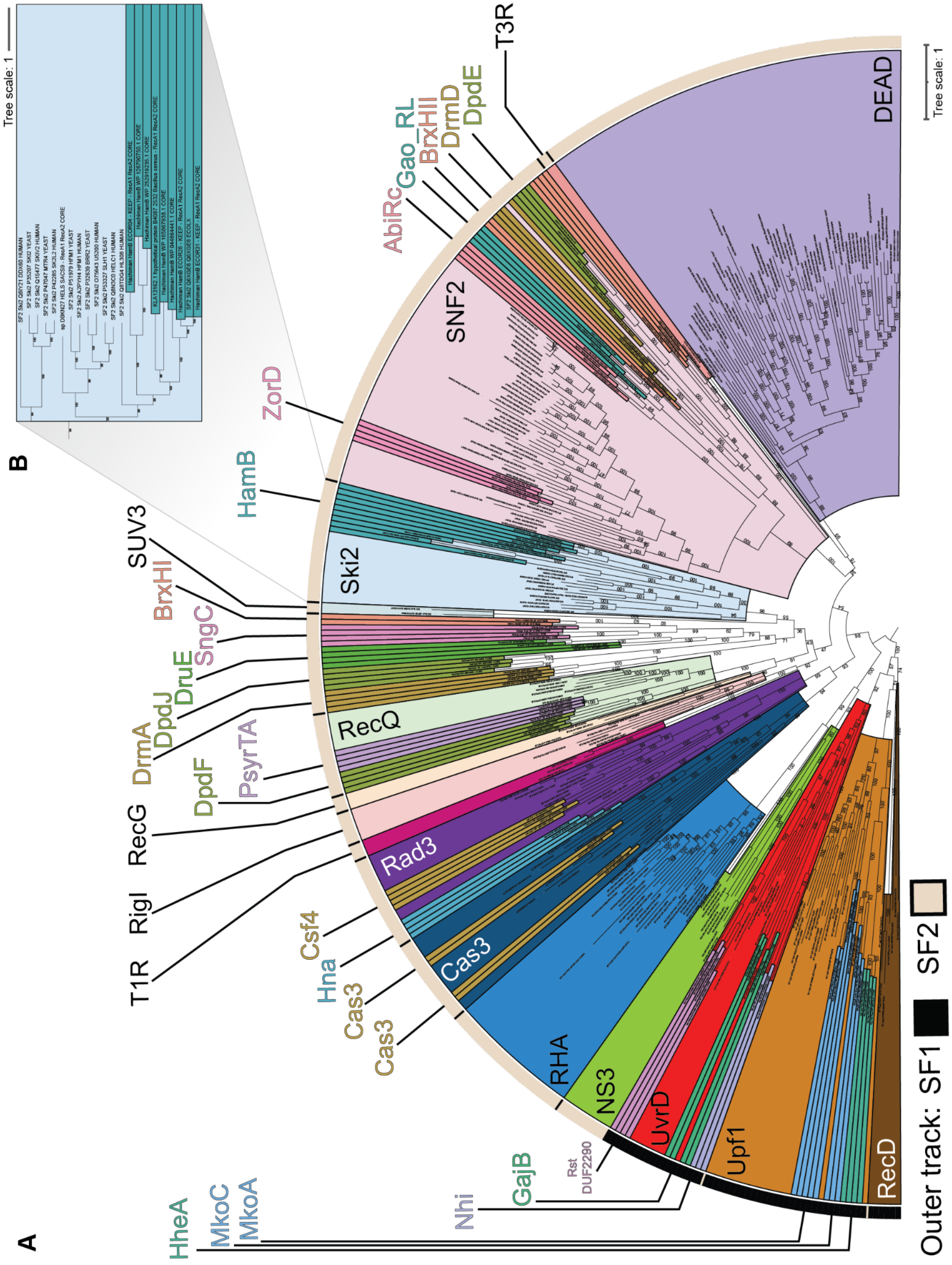


**Figure S1 | Helicases and Hachiman phylogenetic analysis. (A)** Annotated phylogenetic tree of phage defense system associated helicase core domains and reference helicases shown in Fig. 1B. Bootstrapping values determined by UFBoot2[72] are shown. **(B)** Zoomed-in view focusing on represented Ski2 helicases.

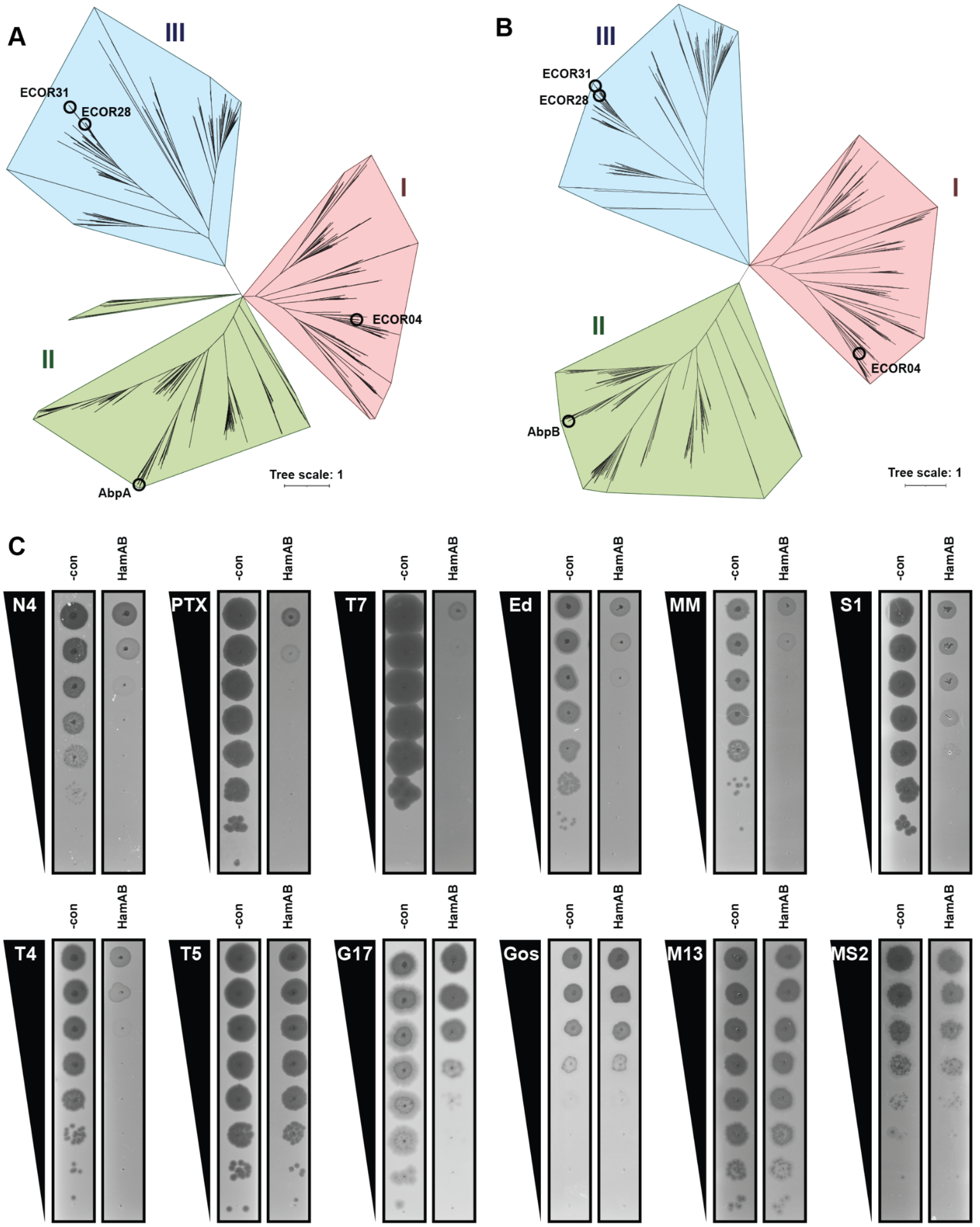


**Figure S2 | Function and phylogeny of tested HamAB proteins.** **(A, B)** Phylogenetic tree of (A) HamA and (B) HamB from DefenseFinder[17] with HamA and HamB sequences from this manuscript and AbpAB[34] are labeled. We assign 3 potential clades of HamB and their corresponding HamA clades as I-III. **(C)** Representative plaque assays for ECOR31 HamAB and phages tested in this study.

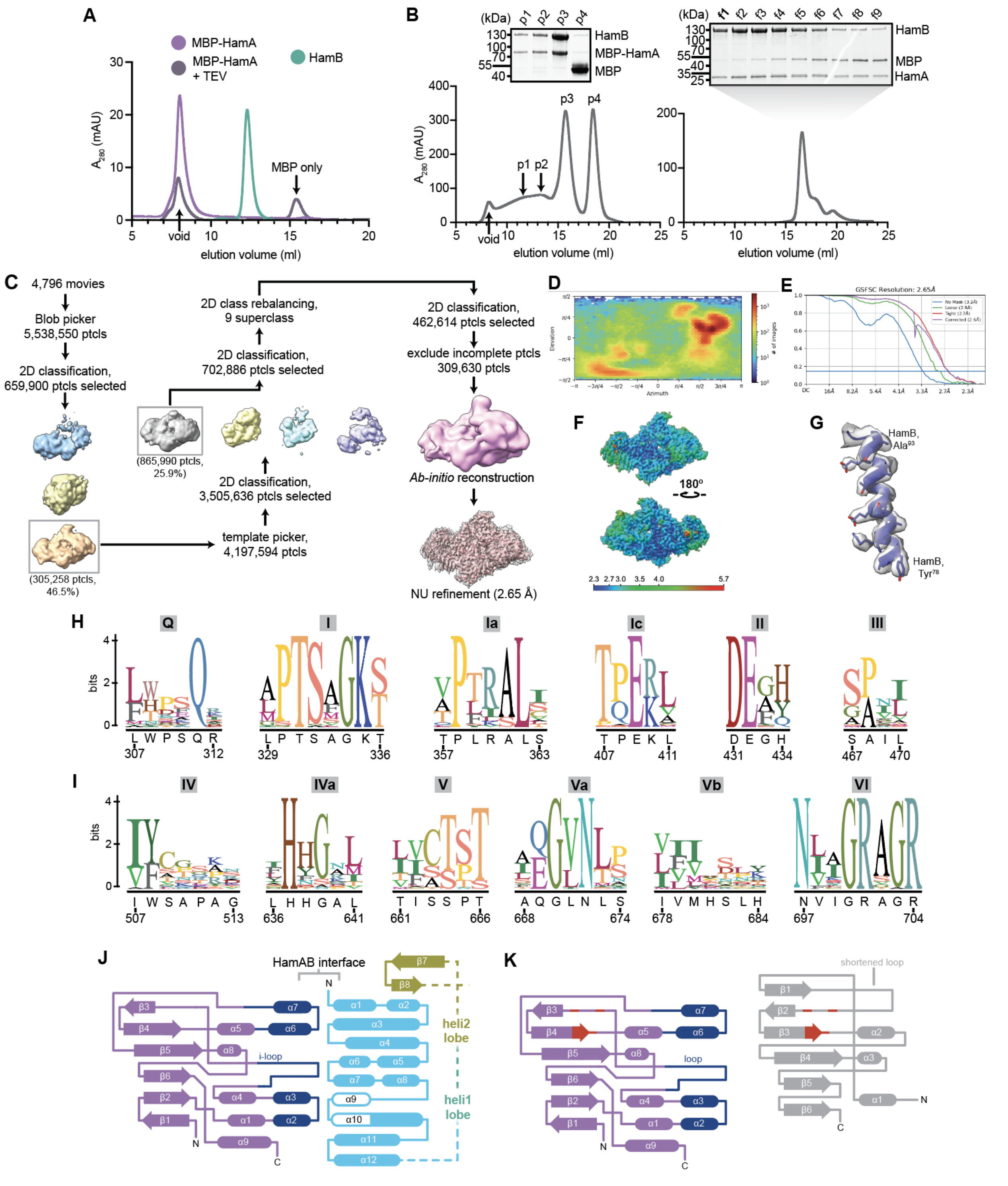


**Figure S3 | Cryo-EM structure of the HamAB apo complex.** **(A)** Size exclusion chromatography traces of MBP-HamA pre-TEV treatment, post-TEV treatment and HamB alone. **(B)** Left, size exclusion chromatography trace of MBP-HamAB and corresponding peaks run on a coomassie PAGE gel. Right, size exclusion chromatography trace of HamAB after TEV protease treatment, with elution fractions run on coomassie PAGE gel shown above. **(C)** Particle picking, classification, and refinement strategy to generate the final apo HamAB density. **(D)** Orientation distribution of the final particle set. **(E)** Gold standard FSC curve. **(F)** Sharpened map colored by local resolution. **(G)** Example model-to-map fit. **(H-I)** Sequence logos of helicase motifs in the RecA1 (H) and RecA2 (I) domains calculated from the HamB MSA. The residue number and identity of the corresponding sequence in ECOR31 HamB is shown below start and end motif positions. **(J)** Secondary structure diagram depicting the HamAB interaction interface. **(K)** Comparison of secondary structures of HamA and the *P. aquatile* Type IIS restriction endonuclease.

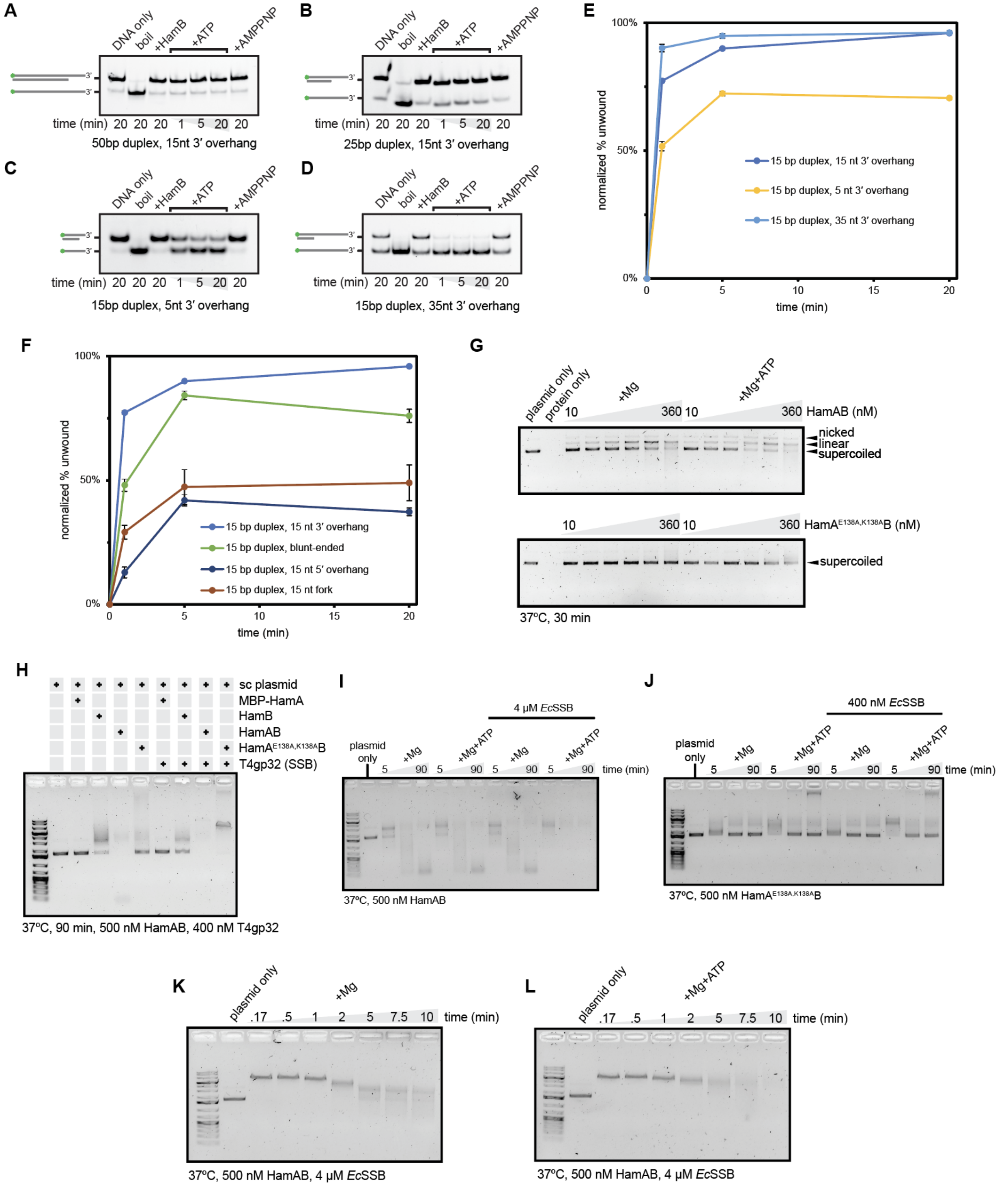


**Figure S4 | Analysis of HamAB biochemical activities *in vitro*. (A-D)** HamB DNA unwinding assays on substrates with a 50 bp duplex and a 15 nt 3′ OH (A), 15 bp duplex and 15 nt 3′ OH (B), 15 nt duplex with a 5 nt 3′ OH (D), and 15 nt duplex with a 35 nt 3′ OH (E). DNA substrates are labeled with 5′ FAM. Gels are representative of three independent biological replicates. **(E-F)** Normalized percent unwinding of DNA substrates with varying OH lengths (E) and different OHs (F). All substrates are labeled with 5′ FAM. Shown are the mean and standard error of quantifications of three replications of unwinding assays in the format of A-D normalized against basal unwinding (see Methods). **(G)** Titration of HamAB WT and HamA*B complexes with plasmid DNA. **(H)** Comparison of Hachiman protein and complex plasmid activities with or without phage T4 SSB. **(I)** Time course of plasmid degradation by HamAB with or without ATP and saturating amounts of *E. coli* SSB. **(J)** Time course of plasmid degradation by HamA*B with or without ATP and limiting amounts of *E. coli* SSB. Conditions in absence of *E. coli* SSB are also shown in Fig. 3H. **(K)** Plasmid degradation over 10 mins with saturating *E. coli* SSB in the absence of ATP. **(L)** Plasmid degradation over 10 mins with saturating *E. coli* SSB and ATP supplied.

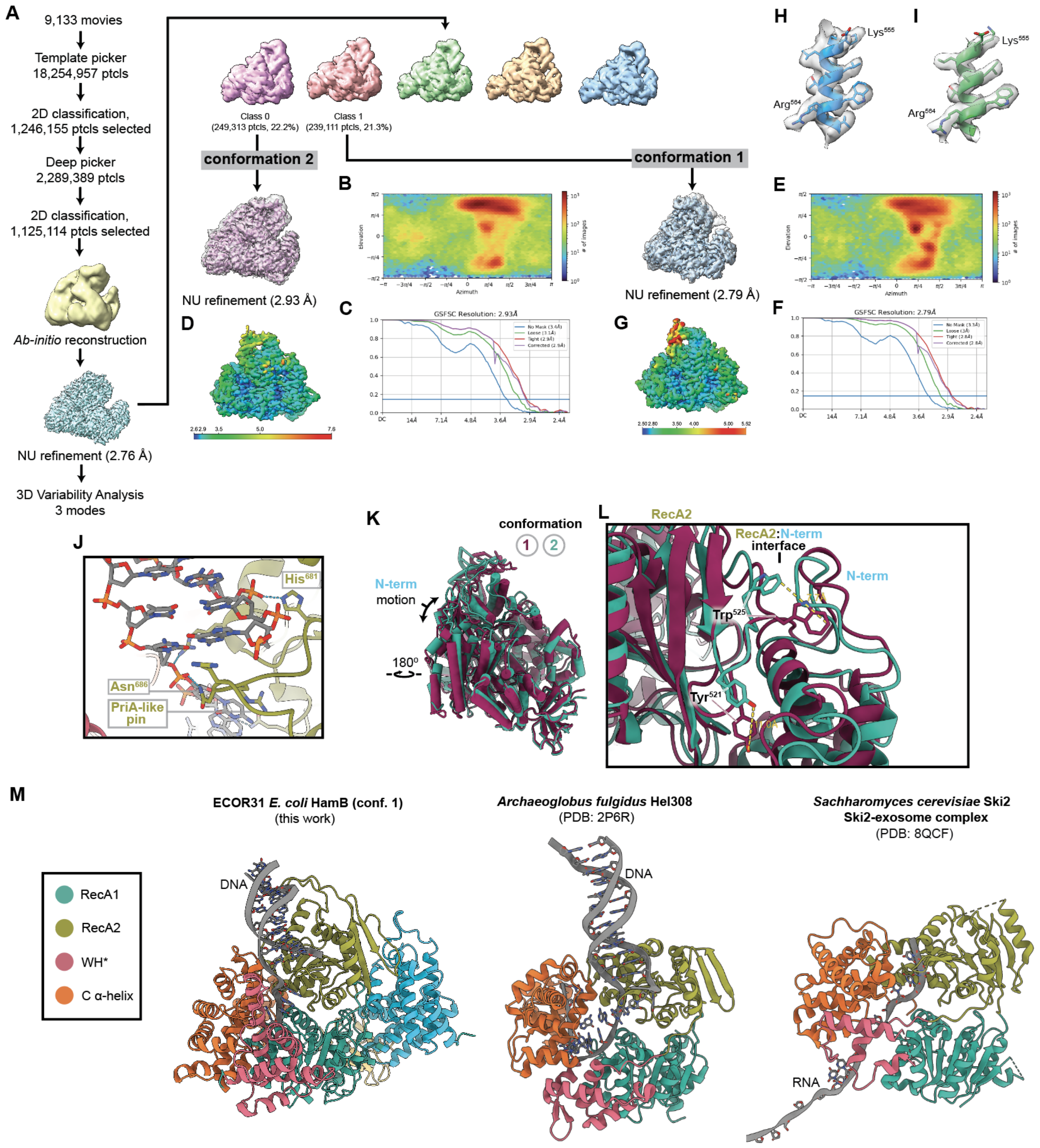


**Figure S5 l Cryo-EM structure of two HamB-DNA complex conformations.** **(A)** Particle picking, classification, and refinement strategy to generate the final HamB-DNA densities for conformations 1 and 2. **(B)** Orientation distribution of the final conformation 2 particle set. **(C)** Gold standard FSC curve for conformation 2. **(D)** Conformation 2 Sharpened map colored by local resolution. **(E)** Orientation distribution of the final conformation 1 particle set. **(F)** Gold standard FSC curve for conformation 1. **(G)** Conformation 1 sharpened map colored by local resolution. **(H)** Example model-to-map fit for conformation 2. **(I)** Example model-to-map fit for conformation 1. **(J)** Molecular detail of the PriA-like strand unwinding pin in HamB-DNA conformation 1. **(K)** Superimposition of conformations 1 and 2 viewed from the HamA side. **(L)** Detail of the HamB RecA2-NAH interface and comparison of conformational changes. **(M)** Comparison of HamB-DNA with other related helicases bound to their substrates. Middle, *A. flugidus* Hel308 (PDB: 2P6R). Right, Ski2-RNA from a structure of the *S. cerevisiae* Ski2-exosome complex (PDB: 8QCF). The exosome complex was hidden for clarity.

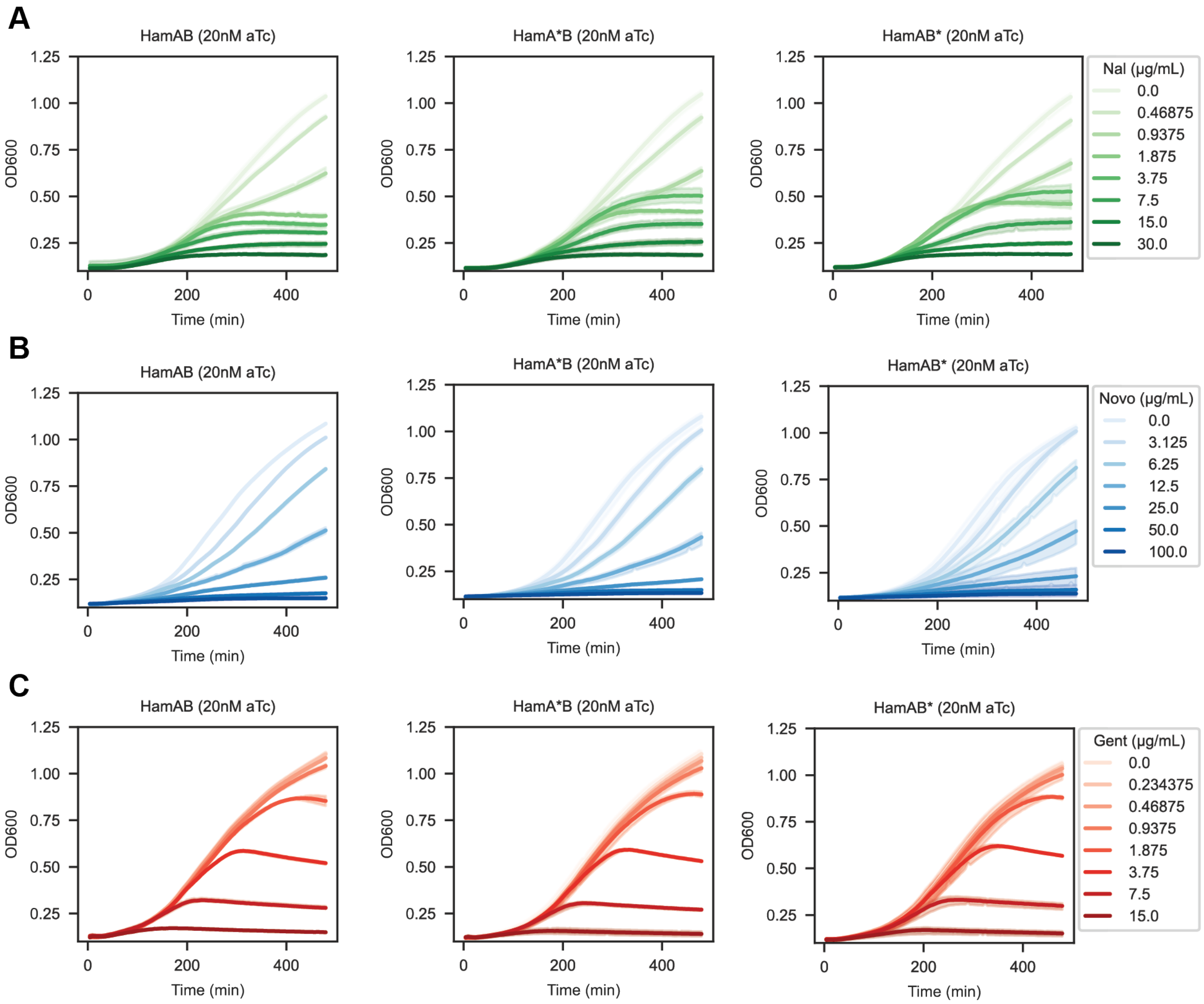


**Figure S6 | Drug-induced DNA damage activates Hachiman. (A-C)** Cell growth of *E. coli* expressing wildtype HamAB (left), nuclease-deficient HamA*B (middle) and helicase-deficient HamAB* (right) at 20nM aTc in the absence or presence of subinhibitory and inhibitory concentrations of nalidixic acid (A), novobiocin (B) or gentamycin (C). Growth curves are colored according to condition. All growth curves performed in biological triplicate.

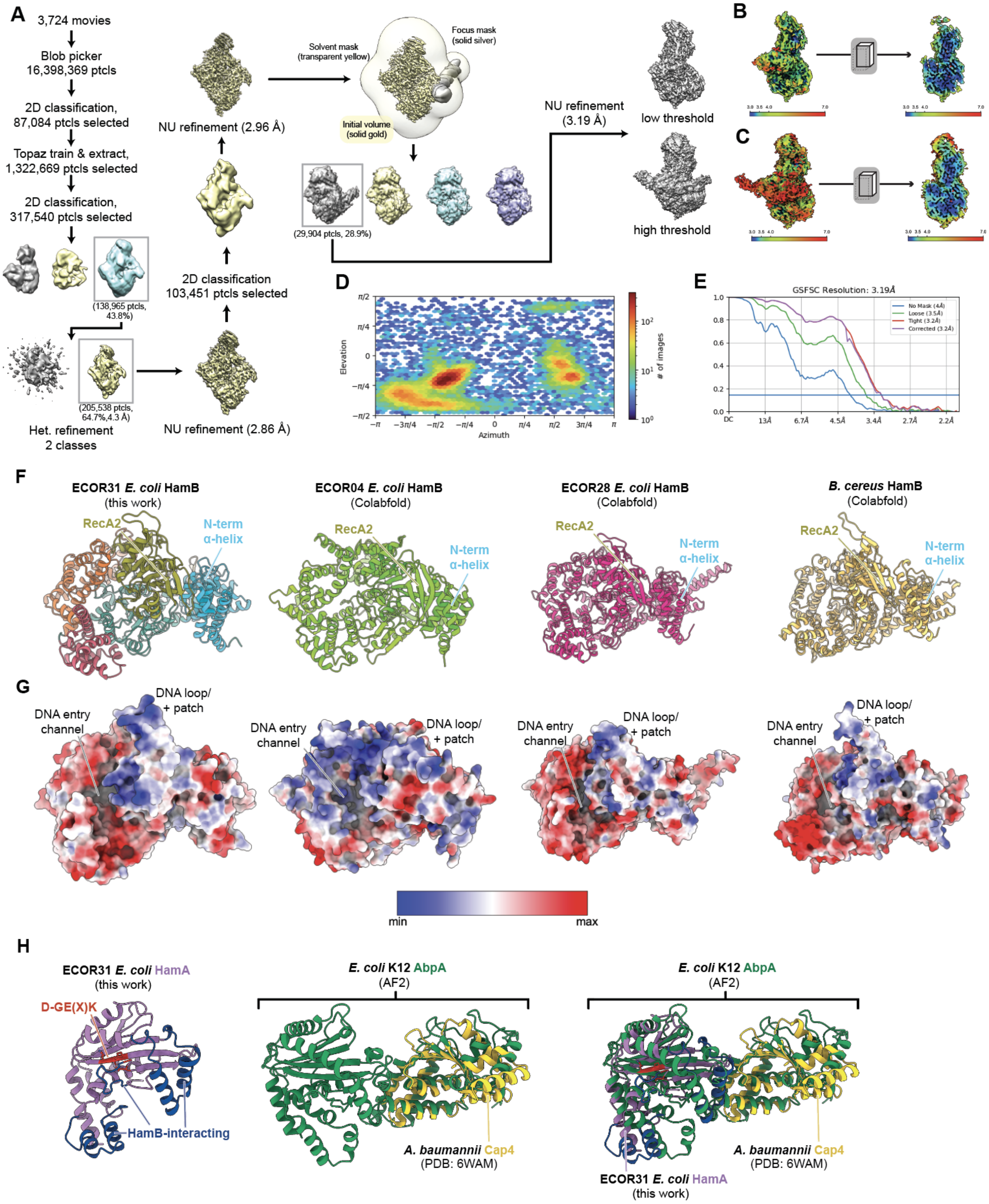


**Figure S7 I Cryo-EM structure of a HamA(E138A, K140A)B-plasmid DNA complex.** **(A)** Particle picking, classification, and refinement strategy to generate the final HamAE138A, K140AB-plasmid DNA density. **(B-C)** Final densities at low (B) and high (C) thresholds colored by local resolution. An inner surface slice is shown to the right. **(D)** Orientation distribution of the final particle set. **(E)** Gold-standard FSC curve. **(D)** Conformation 2 sharpened map colored by local resolution. **(E)** Orientation distribution of the final conformation 1 particle set. **(F)** Comparison of ECOR31 HamB from this study with ECOR04 HamB, ECOR28 HamB, and *Bacillus cereus* HamB Colabfold predictions. **(G)** Electrostatic surface potential representations of the structures from F, in the same orientation and scale. The DNA entry channel and RecA2 DNA loop/ positively charged patch are indicated. **(H)** HamA and Cap4 structural superimpositions with AbpA.

**Table S1 | Cryo-EM data collection, refinement, and validation statistics.**

| | ***E. coli*** **ECOR31 apo HamAB** (EMD-43613, PDB 8VX9) | ***E. coli*** **ECOR31 HamB-DNA (conformation 1)** (EMD-43615, PDB 8VXA) | ***E. coli*** **ECOR31 HamB-DNA (conformation 2)** (EMD-43616, PDB 8VXC) | ***E. coli ECOR31*** **HamA(E138A,K140A)B-plasmid DNA** (EMD-43643, PDB 8VXY) |
|---|---|---|---|---|
| **Data collection and processing** | | | | |
| Voltage (kV) | 300 | 200 | | 300 |
| Electron exposure ($e^-$/Å$^{-2}$) | 50 | 50 | | 50 |
| Defocus range (μm) | -0.8 to -1.8 | -0.8 to -1.8 | | -0.8 to -1.8 |
| Pixel size (Å) | 1.05 | 1.12 | | 1.05 |
| Symmetry imposed | C1 | C1 | | C1 |
| Initial particle images (no.) | 5,538,550 | 18,254,957 | | 16,398,369 |
| Final particle images (no.) | 309,630 | 239,111 | 249,313 | 29,904 |
| Map Resolution (Å) | 2.65 | 2.79 | 2.93 | 3.19 |
| FSC threshold | 0.143 | 0.143 | 0.143 | 0.143 |
| Map resolution range (Å) | 2.3-5.7 | 2.5-4.8 | 2.6-7.6 | 2.9-7.0 |
| **Refinement** | | | | |
| Initial model used | AF2 model | AF2 model | 8VXA (this work) | 8VX9 (this work) |
| Model resolution (Å) | 3.08 | 3.18 | 3.23 | 3.87 |
| FSC threshold | 0.5 | 0.5 | 0.5 | 0.5 |
| Map-sharpening B-factor (Å$^2$) | -85.9 | -92.5 | -93.9 | -60.9 |
| Model composition | | | | |
| Non-hydrogen atoms | 11,244 | 9,431 | 9,391 | 12,324 |
| Protein residues | 1,433 | 1,139 | 1,139 | 1,406 |
| Nucleotide residues | 0 | 24 | 22 | 62 |
| Ligands | 0 | 0 | 0 | 1 |
| B-factor (Å$^2$) | | | | |
| Proteins | 50.88 | 53.27 | 51.88 | 111.72 |
| Nucleic acid | - | 83.34 | 91.38 | 314.69 |
| Ligands | - | - | - | 89.85 |
| R.M.S. deviations | | | | |
| Bond lengths (Å) | 0.27 | 0.003 | 0.003 | 0.003 |
| Bond angles (°) | 0.52 | 0.530 | 0.506 | 0.571 |
| **Validation** | | | | |
| MolProbity score | 1.87 | 1.73 | 1.68 | 1.99 |
| Clashscore | 11.85 | 10.67 | 9.14 | 14.11 |
| Poor rotamers (%) | 0.08 | 0 | 0 | 0 |
| Ramachandran plot | | | | |
| Favored (%) | 95.94 | 96.92 | 96.83 | 95.21 |
| Allowed (%) | 4.06 | 3.08 | 3.17 | 4.79 |
| Disallowed (%) | 0 | 0 | 0 | 0 |

**Table S2 I Strains used in this study**

| Strain | Plasmid | Use |
|---|---|---|
| *E.coli* dh10b | N/A | Phage assays, antibiotic sensitivity assays, microscopy, cloning |
| *E.coli* BL21-AI | N/A | Protein expression and purification |
| *E.coli* BW25113 | N/A | Phage propagation |
| *E.coli* dh5a F' | F' | Propagation and assaying phages MS2 and M13 |
| *E.coli* DSM103255 | N/A | Propagation of phage G17 |
| *E.coli* MC1000 | N/A | Propagation and assaying phage Goslar |
| *E.coli* ECOR47 | N/A | Assaying phage G17 |
| *E.coli* ECOR04 | N/A | Wildtype locus of ECOR04 Hachiman |
| *E.coli* ECOR28 | N/A | Wildtype locus of ECOR28 Hachiman |
| *E.coli* ECOR31 | N/A | Wildtype locus of ECOR31 Hachiman |

**Table S3 | Plasmids used in this study**

| Plasmid Number | Description | Origin | Resistance | Use | Source |
|---|---|---|---|---|---|
| pBA635_ddCasRX_RFP_CDS_1 | Catalytically deactivated RfxCas13d under pTet control (using RFP-targeting guide) | p15a | Cm | Negative control for plaque assays. | [88] |
| pBA1368_pTet-ECOR04_Hachiman | Hachiman from ECOR04 under pTet control. | p15a | Cm | Bacterial assays for ECOR04 Hachiman including phage infection assays. | Current study |
| pBA1369_pTet-ECOR28_Hachiman | Hachiman from ECOR28 under pTet control. | p15a | Cm | Bacterial assays for ECOR28 Hachiman including phage infection assays. | Current study |
| pBA1370_pTet-ECOR31_Hachiman | Hachiman from ECOR31 under pTet control. | p15a | Cm | Bacterial assays for ECOR31 Hachiman including phage infection assays and antibiotic sensitivity assays. | Current study |
| pBA1444_ECOR31HamA_purification | MBP-His-tagged ECOR31 HamA for protein purification | ColE1 | Carb | Protein expression and purification of ECOR31 HamA. | Current study |
| pBA1445_ECOR31HamB_purification | MBP-His-tagged ECOR31 HamB for protein purification | ColE1 | Carb | Protein expression and purification of ECOR31 HamB. | Current study |
| pBA1464_pTet-ECOR31Ham_DELhamA_Mut | Hachiman from ECOR31 under pTet control with HamA deleted (and HamB native RBS intact). | p15a | Cm | Bacterial assays for mutant ECOR31 Hachiman including phage infection assays. Mutant encodes for a HamA deletion, leaving the HamB RBS intact. | Current study |
| pBA1465_pTet-ECOR31Ham_DELhamB_Mut | Hachiman from ECOR31 under pTet control with HamB deleted. | p15a | Cm | Bacterial assays for mutant ECOR31 Hachiman including phage infection assays. Mutant encodes for a HamB deletion. | Current study |
| pBA1467_pTet-ECOR31Ham_hamB_walkerB_Mut | Hachiman from ECOR31 under pTet control with HamB mutated (D431A). | p15a | Cm | Bacterial assays for mutant ECOR31 Hachiman including phage infection assays and antibiotic sensitivity assays. Mutant encodes for a HamB helicase active site mutant. | Current study |
| pBA1468_pTet-ECOR31Ham_hamA_D119A_Mut | Hachiman from ECOR31 under pTet control with HamA mutated (D119A). | p15a | Cm | Bacterial assays for mutant ECOR31 Hachiman including phage infection assays. Mutant encodes for a HamA nuclease active site mutant. | Current study |
| pBA1469_pTet-ECOR31Ham_hamA_E138A_K140A_Mut | Hachiman from ECOR31 under pTet control with HamA mutated (E138AK140A). | p15a | Cm | Bacterial assays for mutant ECOR31 Hachiman including phage infection assays and antibiotic sensitivity assays. Mutant encodes for a HamA nuclease active site double mutant. | Current study |
| pBA1580_ECOR31HamAB_CoPurification | MBP-His-tagged ECOR31 HamA operonic with HamB for protein purification | ColE1 | Carb | Protein expression and purification of ECOR31 HamAB complex. | Current study |
| pBA1583_ECOR31HamA(E138AK140A)B_CoPurification | MBP-His-tagged ECOR31 mutant HamA (E138AK140A) operonic with HamB for protein purification | ColE1 | Carb | Protein expression and purification of mutant ECOR31 HamAB complex. Mutant encodes for a HamA nuclease active site mutant. Mutant encodes for a HamA nuclease active site double mutant. | Current study |
| pBA1606_pTet-ECOR31HamAB_Mut_HamAΔdza 166-195 | Hachiman from ECOR31 under pTet control with HamA mutated (Δ166-195). | p15a | Cm | Bacterial assays for mutant ECOR31 Hachiman including phage infection assays. Mutant encodes for a HamAB interface mutant in HamA. | Current study |
| pBA1607_pTet-ECOR31HamAB_Mut_HamA_mut3 RXXK_AXXA (104, 108) | Hachiman from ECOR31 under pTet control with HamA mutated (R104A, K108A). | p15a | Cm | Bacterial assays for mutant ECOR31 Hachiman including phage infection assays. Mutant encodes for a HamAB interface mutant in HamA. | Current study |

**Table S4 l Phages used in this study**

| Phage | Propagation Host | Assay Host | Class (ICTV 2022) | Genus (ICTV 2022) | Culture Notes |
|---|---|---|---|---|---|
| EdH4 | *E.coli* BW25113 | *E.coli* BW25113 | Caudoviricetes | Vequintavirus | |
| G17 | DSM103255 | *E.coli* ECOR47 | Caudoviricetes | Asteriusvirus | Plaque assays require 0.375% top agar |
| Goslar | *E.coli* MC1000 | *E.coli* MC1000 | Caudoviricetes | Goslarvirus | Plaque assays require 0.375% top agar |
| M13 | *E.coli* Dh5a F' | *E.coli* Dh5a F' | Faserviricetes | Inovirus | Add 1mM CaCl2, Need F' plasmid |
| MM02 | *E.coli* BW25113 | *E.coli* BW25113 | Caudoviricetes | Mosigvirus | |
| MS2 | *E.coli* Dh5a F' | *E.coli* Dh5a F' | Leviviricetes | Emesvirus | Add 1mM CaCl2, Need F' plasmid |
| N4 | *E.coli* BW25113 | *E.coli* BW25113 | Caudoviricetes | Enquatrovirus | |
| PTXU 04 | *E.coli* BW25113 | *E.coli* BW25113 | Caudoviricetes | Xuquatrovirus | |
| SUSP 1 | *E.coli* BW25113 | *E.coli* BW25113 | Caudoviricetes | Suspvirus | |
| T4 | *E.coli* BW25113 | *E.coli* BW25113 | Caudoviricetes | Tequatrovirus | |
| T5 | *E.coli* BW25113 | *E.coli* BW25113 | Caudoviricetes | Tequintavirus | |
| T7 | *E.coli* BW25113 | *E.coli* BW25113 | Caudoviricetes | Teseptimavirus | |

**Table S5 | ATPase assay oligonucleotide substrates**

| Number | Substrate Description in Figure 3A | Sequence (5' to 3', r prefix represents RNA) | sequence length (nt) |
|---|---|---|---|
| 1 | polyT ssDNA | TTTTTTTTTTTTTTTTTTTTTTTTTTTTTTTTTTTTTTTT | 40 |
| 2 | mixed ssDNA | TCGTCACCAGTACAAACTACAACGCCTGTAGCATTCCACA | 40 |
| 3 | 5' overhang DNA | TCGTCACCAGTACAAAC | 17 |
| | | TTTTTTTTTTTTTTTTGTTTGTACTGGTGACGA | 33 |
| 4 | 3' overhang DNA | TCGTCACCAGTACAAAC | 17 |
| | | GTTTGTACTGGTGACGATTTTTTTTTTTTTTTT | 33 |
| 5 | complete duplex DNA | TCGTCACCAGTACAAAC | 17 |
| | | GTTTGTACTGGTGACGA | 17 |
| 6 | polyA ssRNA | rA-<br>rArArArArArArArArArArArArArArArArArArArArA<br>rArArArArArArArArArArArArArArArArArA | 40 |
| 7 | mixed ssRNA | rUrCrGrUrCrArCrCrArGrUrArCrArArArCrU-<br>rArCrArArCrGrCrCrUrGrU-<br>rArGrCrArUrUrCrCrArCrA | 40 |
| 8 | 5' overhang RNA | rUrCrGrUrCrArCrCrArGrUrArCrArArArC | 17 |
| | | rU-<br>rUrUrUrUrUrUrUrUrUrUrUrUrUrUrUrGrUrUrUrGrU<br>rArCrUrGrGrUrGrArCrGrA | 33 |
| 9 | 3' overhang RNA | rUrCrGrUrCrArCrCrArGrUrArCrArArArC | 17 |
| | | rGrUrUrUrGrUrArCrUrGrGrUrGrArCr-<br>GrArUrUrUrUrUrUrUrUrUrUrUrUrUrUrUrU | 33 |
| 10 | dsRNA | rUrCrGrUrCrArCrCrArGrUrArCrArArArC | 17 |
| | | rGrUrUrUrGrUrArCrUrGrGrUrGrArCrGrA | 17 |
| 11 | RNA/DNA hybrid w/ 3' DNA overhang | GTTTGTACTGGTGACGATTTTTTTTTTTTTTTT | 33 |
| | | rUrCrGrUrCrArCrCrArGrUrArCrArArArC | 17 |
| 12 | DNA/RNA hybrid w/ 3' RNA overhang | TCGTCACCAGTACAAAC | 17 |
| | | rGrUrUrUrGrUrArCrUrGrGrUrGrArCr-<br>GrArUrUrUrUrUrUrUrUrUrUrUrUrUrUrUrU | 33 |
| 13 | blunt dsDNA/RNA hybrid | TCGTCACCAGTACAAAC | 17 |
| | | rGrUrUrUrGrUrArCrUrGrGrUrGrArCrGrA | 17 |

**Table S6 | Unwinding assay oligonucleotide substrates**

| Number | Substrate Description | Sequence (5' to 3', * represents 6-FAM) | sequence length (nt) |
|---|---|---|---|
| 1 | 15bp duplex, with 15nt 3' overhang | *ATATCGTAGGTATGGTGGAGGCCGGTAGGT | 30 |
| | | CCATACCTACGATAT | 15 |
| 2 | 25bp duplex, with 15nt 3' overhang | *ATATCGTAGGTATGGTGGAGGCCGG-TAGGTAATTCTAGCC | 40 |
| | | CCGGCCTCCACCATACCTACGATAT | 25 |
| 3 | 50bp duplex, with 15nt 3' overhang | *ATATCGTAGGTATGGTGGAGGCCGG-TAGGTAATTCTAGCCCAGCTCCATGCGAGCACTAC-CAATC | 65 |
| | | CATGGAGCTGGGCTAGAATTACCTACCGGCCTCCAC-CATACCTACGATAT | 50 |
| 4 | 15bp duplex, with 5nt 3' overhang | *ATATCGTAGGTATGGTGGAG | 20 |
| | | CCATACCTACGATAT | 15 |
| 5 | 15bp duplex, with 35nt 3' overhang | *ATATCGTAGGTATGGTGGAGGCCGG-TAGGTAATTCTAGCCCAGCTCGATG | 50 |
| | | CCATACCTACGATAT | 15 |
| 6 | 15bp duplex, blunt-ended | *ATATCGTAGGTATGG | 15 |
| | | CCATACCTACGATAT | 15 |
| 7 | 15bp duplex, with 15nt 5' overhang | *ATATCGTAGGTATGGTGGAGGCCGGTAGGT | 30 |
| | | ACCTACCGGCCTCCA | 15 |
| 8 | 15bp duplex, with 15nt fork | *ATATCGTAGGTATGGATCATTGGCGGATGA | 30 |
| | | GGTCATTAGTCCTCGCCATACCTACGATAT | 30 |
